# Alpha Spatial: Language-guided prediction of single-cell spatial gene expression from histology

**DOI:** 10.64898/2026.10.08.756444

**Authors:** Xi Yang, Ziyi Guo, Xuyuan Guo, Wan Wang, Bowen Bai, Xingran Liu, Chao Zhang, Yufei He, Yuncai Liu

## Abstract

Routine histology preserves tissue architecture but leaves its molecular organization largely unmeasured. Recovering this information requires learning across heterogeneous gene panels and constraining a biologically ill-posed morphology-to-expression mapping. Here we present Alpha Spatial (AS), a language-conditioned framework that predicts cell-level gene expression from histology while jointly learning from different tissues and panels. Matched biological context improved prediction over omitted context, whereas incompatible context reduced accuracy. Multi-tissue learning further improved breast-tissue prediction, and AS outperformed the evaluated methods while preserving molecular-state organization. In breast cancer, AS recovered *POSTN*-associated stromal programmes and HER2-associated molecular and spatial differences. Exploratory patient-level analyses identified a *CXCL14*-associated spatial state whose abundance was associated with non-response to trastuzumab-containing treatment and larger residual invasive tumours. These findings establish a framework for reusing heterogeneous molecular observations to investigate tissue organization and treatment-associated microenvironments in archived histology.

## Introduction

Every archived histological slide preserves a record of how cells were arranged within a patient’s tissue^1–3^. Across large clinical collections, these records offer an opportunity to investigate how tumour microenvironments differ between patients and relate to treatment outcomes^4–8^. Yet their molecular interpretation remains limited: routine haematoxylin and eosin (H&E) staining reveals cellular morphology and tissue architecture, but does not directly measure the gene-expression programmes that distinguish cellular states. Recovering this information from existing slides could extend molecular investigation to retrospective cohorts that were never profiled with spatial assays.

Spatial transcriptomics has made this possibility increasingly tangible^9–11^. By pairing molecular measurements with histological images, computational approaches have demonstrated that tissue morphology contains information predictive of gene expression^12–14^. The remaining challenge, however, extends beyond obtaining high correlations for individual genes. Understanding a microenvironment requires recovering molecular differences between cells while retaining the relationships that organize them into tissue^15–17^. Conventional spot-level measurements combine signals from neighbouring cells^18,19^, whereas cell-resolved targeted assays observe only predefined gene panels that differ across studies^20–23^. An unmeasured gene is not equivalent to a measured gene with no detected transcripts. Joint learning must therefore accommodate differences in molecular coverage without discarding valid panel-specific observations or treating missing measurements as zeros^24^.

These differences raise a second question concerning the interpretation of morphology itself. Histological appearance constrains molecular possibilities, but the mapping from morphology to expression may depend on biological context. We therefore asked how contextual information could constrain this ill-posed prediction problem while enabling joint learning from heterogeneous molecular measurements. Learning across tissues may further help distinguish shared cellular features from context-dependent molecular associations.

Here we introduce Alpha Spatial (AS), a language-conditioned framework that predicts cell-associated gene expression from histology using a multimodal backbone based on Qwen3.5^25^. Each prediction combines an image patch, a target-cell bounding box and text describing the tissue or disease and assay panel. The bounding box identifies the target cell while preserving surrounding tissue as visual context. AS generates named gene–count pairs rather than a fixed-dimensional expression vector, allowing examples with different supervision gene sets to share an output format. Biomedical and pathology domain pretraining precedes supervised learning from paired images and spatial molecular measurements.

We evaluated whether explicit context and broader tissue experience improved molecular prediction and its biological utility. Matched biological context improved cell-level expression correspondence relative to omitted context, while multi-tissue learning improved prediction within breast tissue. At matched spatial resolution, AS outperformed the evaluated methods in expression prediction and recovery of measured molecular states. Biological applications recovered POSTN-associated fibroblast programmes and their epithelial relationships, as well as HER2-associated molecular and spatial differences. In the trastuzumab-response cohort, greater abundance of a CXCL14-associated spatial state was associated with non-response and larger residual invasive tumours. Together, these analyses connect learning from heterogeneous molecular observations with the investigation of tissue organization and treatment-associated microenvironments in routine histology.

## Results

### Development of Alpha Spatial for language-guided spatial transcriptomic inference

Alpha Spatial (AS) is designed to predict cell-associated gene expression from histological images while retaining the spatial coordinates needed to reconstruct tissue organization. Given an H&E image, a target-cell bounding box and a natural-language description of the tissue or disease context and gene panel, the model generates gene–count pairs for the specified cell. The framework combines domain-knowledge pretraining with supervised learning from paired histological and spatial transcriptomic data, using language to describe both the biological context and the requested prediction task (Fig. 1a–c).

**Fig. 1.**
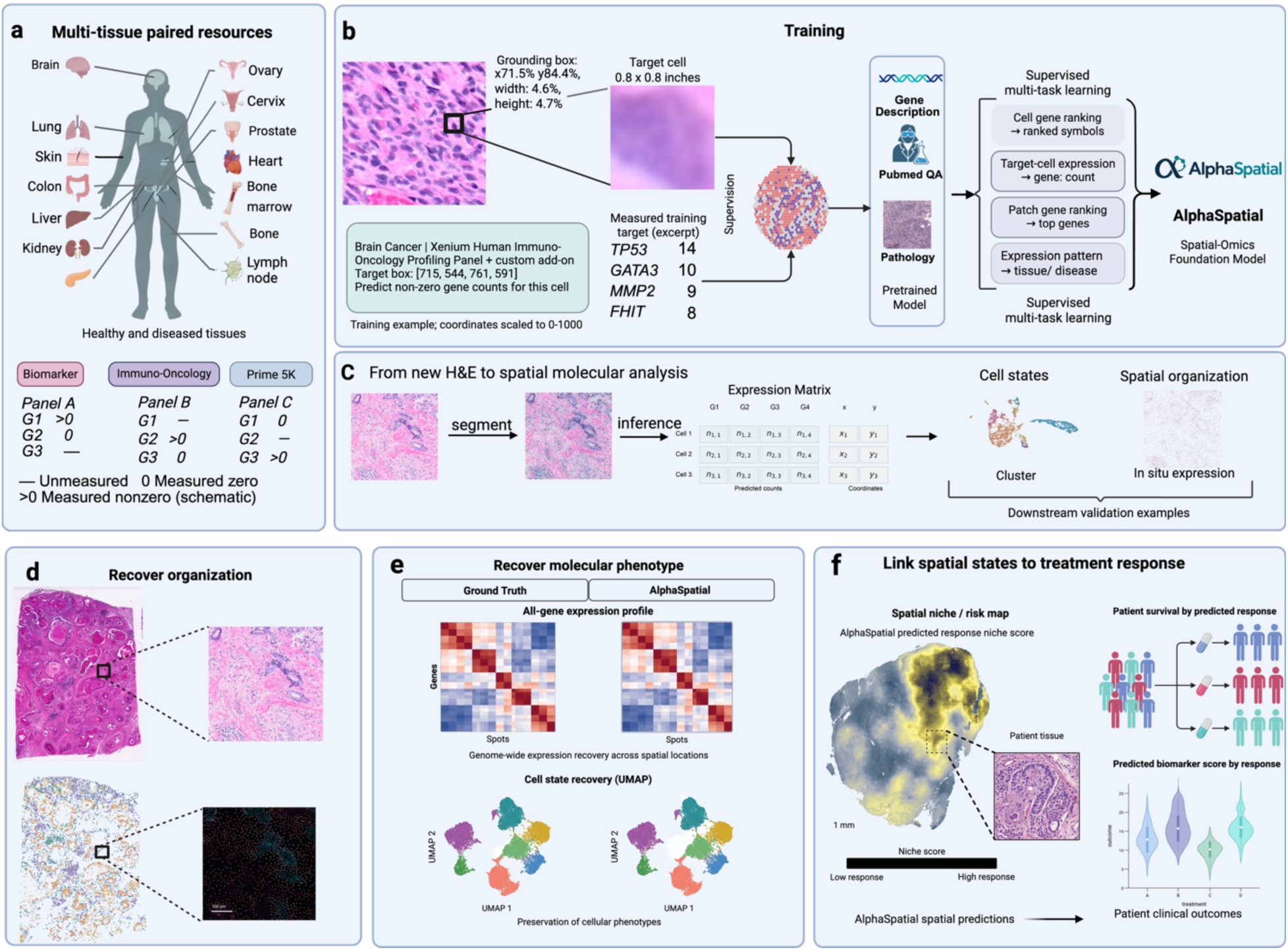
Alpha Spatial framework for language-guided spatial gene-expression predictiona. Paired histological and spatial transcriptomic resources across tissues and gene panels. A dash indicates an unmeasured gene; 0 and >0 indicate measured zero and non-zero counts, respectively. **b** Domain-knowledge pretraining uses gene descriptions, biomedical question–answer pairs and pathology information. Subsequent multimodal training uses paired images and expression, with prompts specifying tissue, panel and target-cell coordinates. Tasks include cell- and patch-level gene ranking, target-cell expression prediction and tissue or disease identification. **c** Cell segmentation and prompt-conditioned inference convert new haematoxylin and eosin (H&E) images into predicted gene counts linked to cell coordinates for molecular and spatial analysis. **d** Spatial organization at tissue and cellular scales. Dashed connectors indicate enlarged regions. **e** Measured and predicted expression profiles and molecular states, visualized using heatmaps and uniform manifold approximation and projection (UMAP). **f** Cohort-level integration of spatial-state distributions and regional burdens with clinical information to investigate associations with treatment response and microenvironment.

We assembled paired resources spanning healthy and diseased tissues profiled using different targeted gene panels (Fig. 1a). These datasets provide complementary molecular observations rather than a uniformly measured expression matrix: a gene may be measured in one dataset but absent from the panel used in another. Importantly, an unmeasured gene is not equivalent to a measured gene with no detected transcripts. We therefore included panel identity alongside tissue or disease information in the training prompts, making the measurement context explicit. Expressing the targets as named gene–count pairs allows training examples with different gene coverage to share a common output format without requiring identical gene lists across datasets.

To provide a biological foundation for learning these image–expression relationships, we first performed domain-knowledge pretraining using gene descriptions, biomedical question–answer pairs and pathology information (Fig. 1b). This stage introduced molecular and histopathological knowledge before supervised training on spatial measurements. The subsequent training stage connected these knowledge resources to paired observations of tissue morphology and gene expression, rather than treating biological descriptions as a substitute for experimentally measured targets.

For spatially grounded supervision, each expression-prediction example combined a histological image with a prompt identifying the target cell by its bounding-box coordinates. This formulation distinguished the cell whose expression was requested while retaining the surrounding tissue as visual context. The corresponding measured expression was represented as non-zero gene–count pairs. We trained the model on four complementary tasks: cell-level gene ranking, target-cell expression prediction, patch-level gene ranking and identification of tissue or disease context from expression patterns (Fig. 1b). These tasks were designed to connect quantitative molecular prediction with expression ordering and biological context at both cellular and regional scales. Natural-language instructions specified the task within the same multimodal framework.

At inference, cell segmentation supplied target-cell coordinates in previously unseen H&E images, and AS predicted expression for individual cells under the specified tissue and panel context (Fig. 1c). The predictions were assembled into a cell-by-gene expression matrix linked to the original spatial coordinates. Paired spatial transcriptomic measurements were therefore used for training and evaluation but were not required as inputs when applying the model to new histological images. This coordinate-linked representation supported both analysis of molecular states across cells and mapping of their distributions within tissue.

We organized the downstream evaluation around three capabilities of this representation. First, we examined whether predicted molecular maps recovered spatial organization across tissue and local cellular scales (Fig. 1d). Second, we compared measured and predicted expression profiles and their associated molecular-state structure, extending evaluation beyond the correlation of individual genes (Fig. 1e). Third, we applied the framework to clinical tissue cohorts to investigate patient-level associations between spatial molecular features and treatment outcomes (Fig. 1f). Together, these analyses assessed whether histology-derived expression could provide a basis for studying molecular heterogeneity, tissue organization and their clinical associations.

### Histology-to-expression prediction is an ill-posed problem without biological context

Histological morphology contains information predictive of gene expression, but whether it sufficiently constrains a cell’s molecular interpretation remains unresolved. We held images and target cells fixed across 79,912 matched cells from eight patients with breast cancer and varied the context between Breast cancer, Breast and Lung cancer. The incompatible Lung cancer prompt reduced mean gene-wise PCC from 0.354 to 0.315, with deterioration in all eight patients (Holm-adjusted p = 0.023). Breast and Breast cancer produced similar overall correlations (0.350 versus 0.354; adjusted p = 0.195; Fig. 2a).

**Fig. 2.**
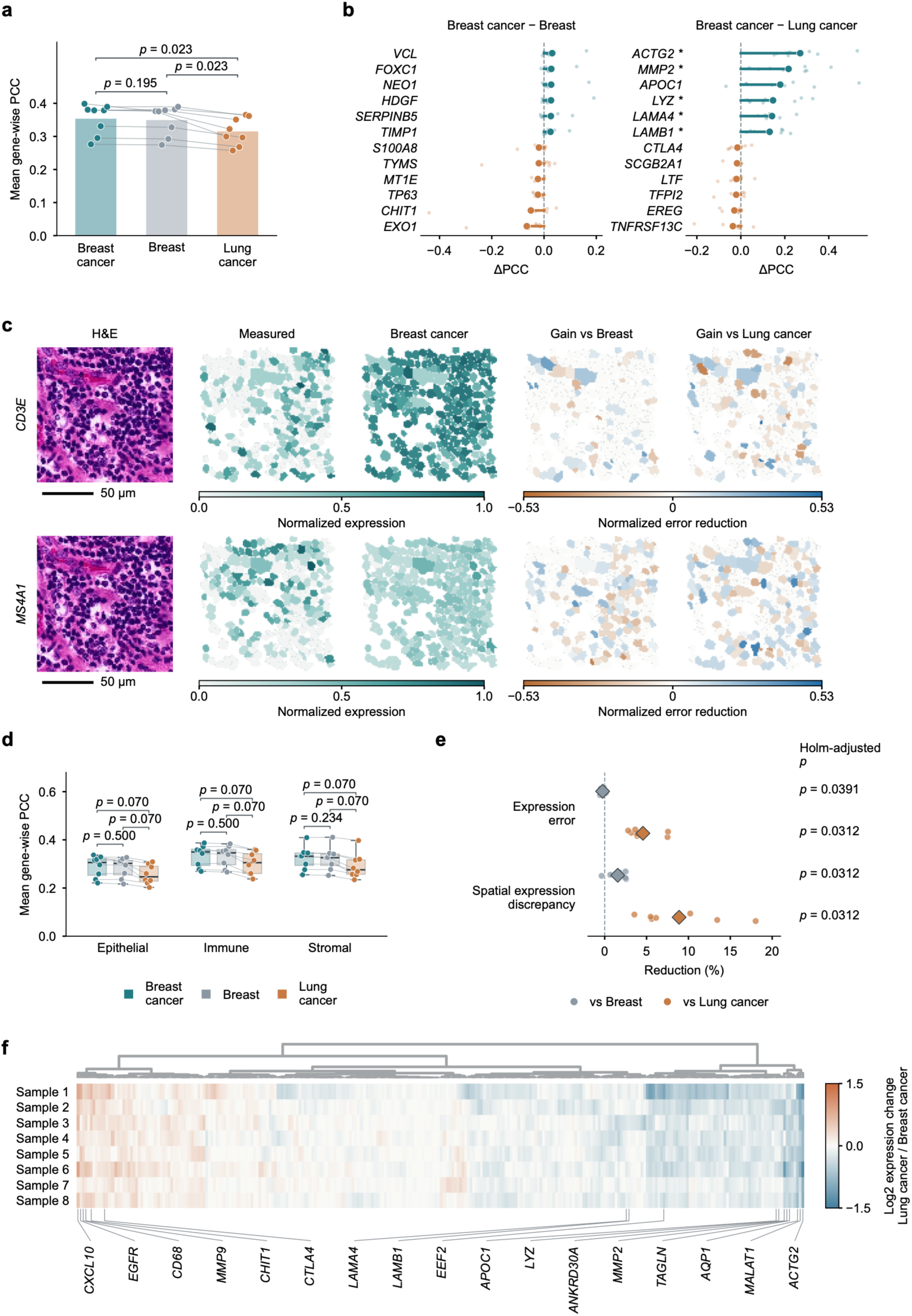
Biological context shapes molecular predictions from fixed histological images. **a,** Mean gene-wise Pearson correlation coefficient (PCC) across eight patients under Breast cancer, Breast and Lung cancer prompts. Bars indicate patient means; connected points denote paired observations. **b,** Genes showing the largest positive and negative mean ΔPCC, calculated as Breast cancer minus the comparator. Large and small points represent means and individual patients, respectively; * indicates Benjamini–Hochberg-adjusted *q* < 0.05. **c,** Representative H&E images, measured and predicted *CD3E* and *MS4A1* expression, and normalized absolute-error reductions. Blue indicates improvement and orange deterioration under the Breast cancer prompt. For each gene, log1p expression is divided by its maximum across measured and all contextual predictions in the displayed field; error reductions use the same scale. Scale bars, 50 μm. **d,** PCC within epithelial, immune and stromal populations annotated from measured expression, excluding annotation genes from evaluation. Boxes indicate medians and interquartile ranges; whiskers extend to 1.5 times the interquartile range. **e,** Relative reductions in log1p expression error and spatial expression discrepancy, measured by root-mean-square error and sliced Wasserstein distance, respectively. Points represent patients and diamonds indicate means. **f,** Changes in patient-level mean predictions across 272 hierarchically clustered genes, expressed as log₂[(Lung cancer + 1)/(Breast cancer + 1)]. Analyses included 79,912 matched cells from eight patients. Tests were two-sided paired Wilcoxon signed-rank tests; Holm adjustment covered three, nine and four comparisons in a, d and e, respectively. Gene-level tests in b were adjusted jointly across genes and both contrasts.

To determine whether correct context provided a benefit beyond avoiding an incompatible prompt, we introduced a condition without tissue, disease or panel information. Across 79,869 four-condition matched cells and identical evaluable genes, Breast cancer increased mean PCC from 0.346 to 0.357 relative to omitted context. This improvement occurred in all eight patients (paired Wilcoxon p = 0.0078). Conversely, Lung cancer reduced PCC to 0.318 (Holm-adjusted p = 0.0391; Supplementary Fig. 1a). Overall expression error did not differ significantly between matched and omitted context (adjusted p = 0.383; Supplementary Fig. 1b). Permuting predicted profiles among cells within each patient preserved each condition’s expression distribution but reduced mean PCC to approximately zero (Supplementary Fig. 1c). Thus, agreement required cell-specific correspondence and could not be explained by the overall molecular background alone.

Importantly, prompt-dependent changes involved genes with shared biological functions. Under the Lung cancer prompt, *ACTA2, ACTG2, TAGLN, MYH11* and *MYLK* decreased in every patient. These genes collectively represent contractile machinery and its regulation: smooth-muscle actins and myosin contribute to force generation, *MYLK* regulates myosin activation, and *TAGLN* participates in actin cytoskeletal organization^26–28^. The coordinated decrease therefore extended across several components of a common functional system. Similarly, *LAMA4, LAMB1* and *COL4A1*, encoding laminin and type IV collagen components of basement membranes, decreased across all eight patients^29^. Conversely, *CD68* and *CHIT1*, previously associated with a shared tumour-associated macrophage differentiation and activation trajectory in breast cancer, increased consistently^30^. These opposing shifts in functionally related gene groups indicate that context changed the biological interpretation assigned to unchanged tissue morphology, rather than merely introducing isolated gene-level errors or uniformly rescaling expression (Fig. 2f).

These functional shifts were accompanied by altered cellular expression patterns and reduced agreement with measurements. Genes with larger downward expression shifts generally exhibited greater PCC losses under the Lung cancer prompt, with a negative relationship in every patient (mean Spearman ρ = −0.49). Changes in non-zero cell fractions showed a corresponding association (mean ρ = −0.39; adjusted p = 0.031; Supplementary Fig. 1). For *ACTG2, MMP2* and *LYZ*, respectively, 46.4%, 21.7% and 12.0% of cells switched between zero and non-zero predictions on average. Together with the spatially heterogeneous errors (Fig. 2b,c), these findings indicate changes in the inferred distribution of expression, rather than uniform rescaling of an otherwise unchanged molecular map.

Providing the matched Breast cancer context reduced expression error by 4.57% and spatial expression discrepancy by 8.89% relative to Lung cancer (both adjusted p = 0.031; Fig. 2e). PCC differences followed the same direction across the three broad cellular compartments, although compartment-level comparisons did not remain significant after adjustment (Fig. 2d). Collectively, these results frame histology-to-expression prediction as a biologically ill-posed inverse problem: the same morphological evidence can receive different, functionally structured molecular interpretations depending on the supplied context. This motivated explicit natural-language conditioning on tissue, disease and measurement panel when learning from heterogeneous spatial transcriptomic resources.

### Multi-tissue learning improves molecular discrimination within breast tissue

We reasoned that learning across tissues could help distinguish molecular features associated with cellular identity from associations specific to a restricted tissue context^31,32^. Broader biological experience could therefore refine predictions within breast tissue, rather than merely extend prediction to additional organs. We compared breast-only and multi-tissue models using 79,980 matched cells from eight patients with breast cancer. Multi-tissue learning increased mean gene-wise Pearson correlation coefficient (PCC) from 0.274 to 0.384 and reduced mean log1p root-mean-square error (RMSE) from 0.711 to 0.535. Both improvements occurred in every patient (Holm-adjusted p = 0.0156 for both comparisons; Fig. 3a).

**Fig. 3.**
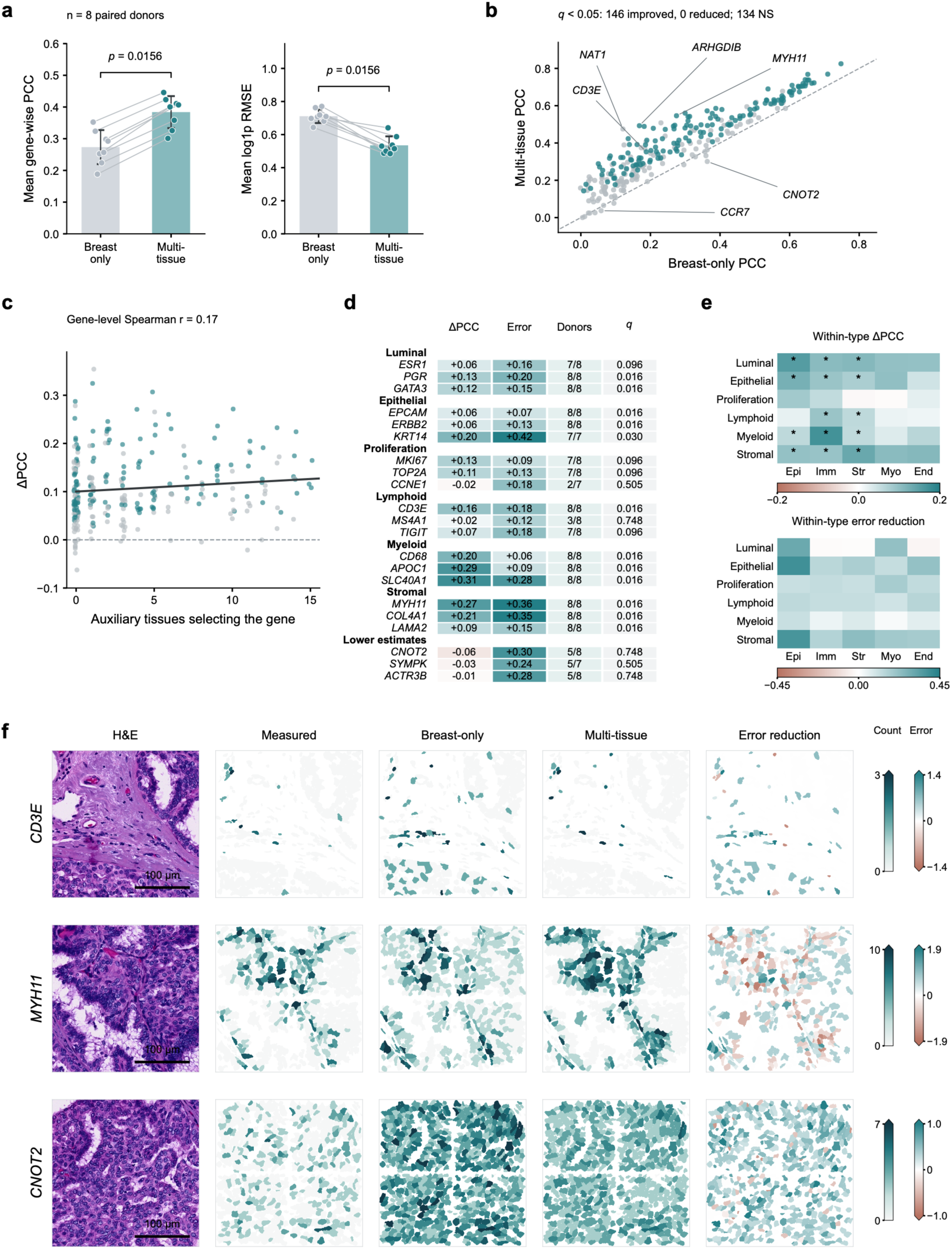
Multi-tissue learning improves molecular prediction within breast tissue. **a,** Mean gene-wise PCC and log1p RMSE across eight paired patients; bars indicate means, error bars s.d., and connected points individual patients. **b,** Patient-averaged PCC for 280 genes. Teal indicates significant improvement (Benjamini–Hochberg-adjusted *q* < 0.05); grey indicates non-significant differences. **c,** Gene-level ΔPCC versus auxiliary-tissue gene-selection breadth, with a descriptive linear fit. **d,** Selected markers and lower-PCC examples. Columns show ΔPCC, error reduction, patients with increased PCC/evaluable patients, and PCC-associated *q* values. **e,** Marker-group performance changes within measured-expression-derived cell annotations (epithelial, immune and stromal: n = 7 each; myoepithelial: n = 4; endothelial: n = 6). Epi, epithelial; Imm, immune; Str, stromal; Myo, myoepithelial; End, endothelial. Asterisks indicate PCC-associated *q* < 0.05; blank entries lack sufficient data. **f,** H&E, measured and predicted expression, and signed absolute-error reduction for representative genes. Colours use log1p counts, with expression ticks labelled in original count units; positive reduction favours multi-tissue prediction. Expression scales are shared within each row and capped at the pooled 99.5th percentile. Scale bars, 100 μm. Tests were two-sided exact patient-paired sign tests, with Holm correction for the two overall endpoints. Gene-level PCC and error tests form separate BH families; within-type tests are corrected separately by metric and cell threshold.

The gains extended across molecular features representing different cellular identities. Across 280 genes, 146 showed increased PCC after false-discovery-rate correction, whereas none showed a significant decrease (Fig. 3b). Improved predictions included epithelial, lymphoid, myeloid and stromal markers, with several gains retained within independently annotated cell populations (Fig. 3d,e). Spatial examples illustrated improved agreement for *CD3E* and *MYH11*, whereas *CNOT2* showed that lower expression error did not necessarily imply higher correlation (Fig. 3f). Improvement was only weakly associated with gene-selection breadth across auxiliary tissues (Spearman ρ = 0.17; Fig. 3c), suggesting that breadth alone did not explain the observed variation.

Further analysis distinguished quantitative accuracy from expression detection. Within measured-positive cells, multi-tissue learning reduced mean log1p RMSE from 0.837 to 0.723 in all eight patients (adjusted p = 0.0313). However, the missed-prediction rate was not reduced, and lower predicted background in measured-zero cells did not reach significance after adjustment (Supplementary Fig. 2a,b). Restricting PCC evaluation to the same evaluable genes preserved the improvement, excluding unequal gene coverage as its explanation (Supplementary Fig. 2c,d). These findings support more accurate quantitative molecular predictions, rather than uniformly improved zero/non-zero discrimination. They are consistent with broader tissue experience refining morphology–expression relationships.

### Alpha Spatial improves the fidelity of spatial expression reconstruction

Having established the benefit of multi-tissue learning, we benchmarked AS against nine histology-to-expression methods: ST-Net^13^, DeepSpaCE^33^, TRIPLEX^34^, EGN v2^35^, BLEEP^36^, STco^37^, mclSTExp^38^, STFlow^39^ and Path2Space^40^. A central consideration was whether a method could learn jointly from valid molecular measurements collected using different gene panels. The inspected native implementations of these comparators used fixed gene spaces and did not directly support joint training across heterogeneous panels while retaining all panel-specific measurements. Such integration requires restricting training to shared genes or modifying the representation and training objective to accommodate unmeasured entries. AS instead makes panel identity explicit, allowing heterogeneous molecular observations to contribute to a shared model without treating unmeasured genes as measured zeros. We therefore evaluated heterogeneous-panel learning as an integral capability of the framework, rather than excluding the multi-tissue model to impose a shared restriction on usable training data.

We evaluated breast-only and multi-tissue AS alongside the comparator methods using specimens from eight patients, a shared 55-µm spatial grid and a fixed 280-gene panel. All methods were assessed at identical evaluable positions and on mutually evaluable genes. Multi-tissue AS achieved a mean gene-wise PCC of 0.430, exceeding both the strongest external comparator, STco (0.303), and breast-only AS (0.290). Its advantage over every comparator was consistent across all eight patients (Benjamini– Hochberg-adjusted q = 0.0099 for each comparison; Fig. 4a,b). This comparison evaluated the complete framework, including its capacity to learn from heterogeneous panels, rather than isolating architecture under identical training data.

**Fig. 4.**
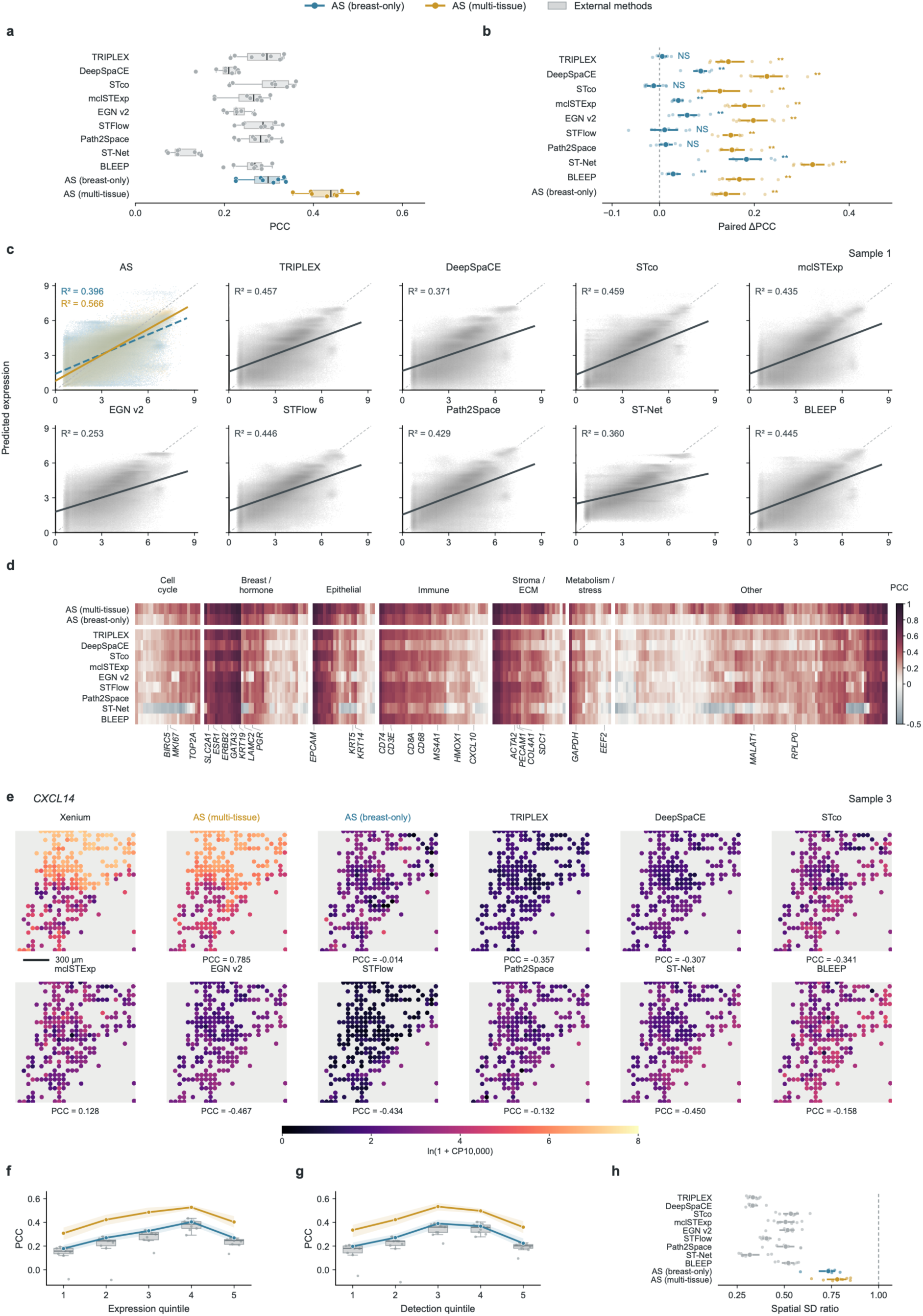
Cross-method evaluation of expression accuracy and spatial variation. **a,** Mean gene-wise spatial PCC across eight patients. **b,** Patient-paired PCC differences between AS and each comparator. **c,** Measured versus predicted expression densities in Sample 1, with linear fits; R² describes pooled spot–gene pairs rather than gene-wise PCC. **d,** Mean PCC for 280 genes, grouped by function and hierarchically ordered within groups. **e,** Selected *CXCL14* expression maps from Sample 3, with local PCC; scale bar, 300 µm. **f,g,** PCC across measured-expression and detection-frequency quintiles. AS curves show patient means; grey boxes summarize the nine external-method means. **h,** Patient-median predicted-to-measured spatial standard-deviation ratios; dashed line, unity. Blue and orange indicate breast-only and multi-tissue AS, respectively. Boxes show medians and interquartile ranges, with whiskers extending to 1.5 times the interquartile range. Intervals and shading indicate 95% patient-bootstrap confidence intervals (20,000 resamples). Expression is ln(1 + CP10,000), calculated within the fixed panel using each prediction’s own total. Comparisons use shared 55-µm positions and mutually evaluable genes. In b, two-sided paired Wilcoxon tests were corrected across 19 comparisons using Benjamini–Hochberg: q < 0.05 *, < 0.01 **, < 0.001 ***; NS, not significant.

The gains extended across biological functions and expression regimes. Gene-level comparisons showed broad improvements across cell-cycle, breast-associated, epithelial, immune, stromal and metabolic gene groups, although individual genes remained variably predictable (Fig. 4d). Multi-tissue AS also achieved the highest mean PCC in every expression and detection-frequency quintile, including low-expression and infrequently detected genes (Fig. 4f,g). Its overall advantage therefore did not depend solely on a small subset of abundant or widely detected transcripts.

Beyond improved correlation, AS better preserved the magnitude and location of expression differences within tissue. In Sample 1, its pooled measured–predicted regression slope was closer to unity (0.743 versus 0.302–0.539 for external methods), with higher R² (0.566 versus 0.253–0.459; Fig. 4c). Across patients, the mean patient-median predicted-to-measured spatial standard-deviation ratio reached 0.780, compared with 0.317–0.535 for external methods, indicating less compression of spatial expression variation (Fig. 4h). The selected *CXCL14* example illustrated the biological consequence of this difference. Multi-tissue AS recovered the regional high–low expression contrast with a local PCC of 0.785, whereas other predictions showed weak or reversed spatial agreement (Fig. 4e). These findings indicate that improved prediction involved both the spatial allocation of expression and the preservation of molecular heterogeneity.

We next examined whether this workflow could operate with target cells identified directly from H&E images. In one breast section, image-derived segmentation objects were matched to 6,191 of 10,000 reference cells using geometry alone. Across 279 evaluable genes, gene-wise PCCs closely tracked those obtained with Xenium-derived localization (R² = 0.973), although mean paired PCC decreased by 0.021 (Supplementary Fig. 3). This single-section comparison supports segmentation-guided molecular prediction from routine histology.

### Alpha Spatial improves prediction accuracy while preserving spatial expression heterogeneity

Accurate prediction of individual genes does not necessarily preserve the molecular relationships that distinguish tissue populations. We therefore examined whether AS reconstructed coordinated expression states and their organization within breast tumours. Across eight patients, measured and predicted profiles were compared at 26,068 matched 55-µm spatial positions using the same 280 genes. For visualization, predictions were projected into a UMAP reference fitted separately to each patient’s measured expression^41^. In Sample 1, AS retained a more distributed representation of the measured states, whereas several comparator predictions concentrated into restricted regions of the reference space (Fig. 5a,b). Extending this comparison to all eight patients showed that this distinction was not confined to the illustrative specimen. AS more closely retained the broad distribution of measured states, whereas comparator projections frequently contracted into narrower regions or showed increased mixing of reference states, particularly in Samples 3, 6 and 7 (Supplementary Fig. 4).

**Fig. 5.**
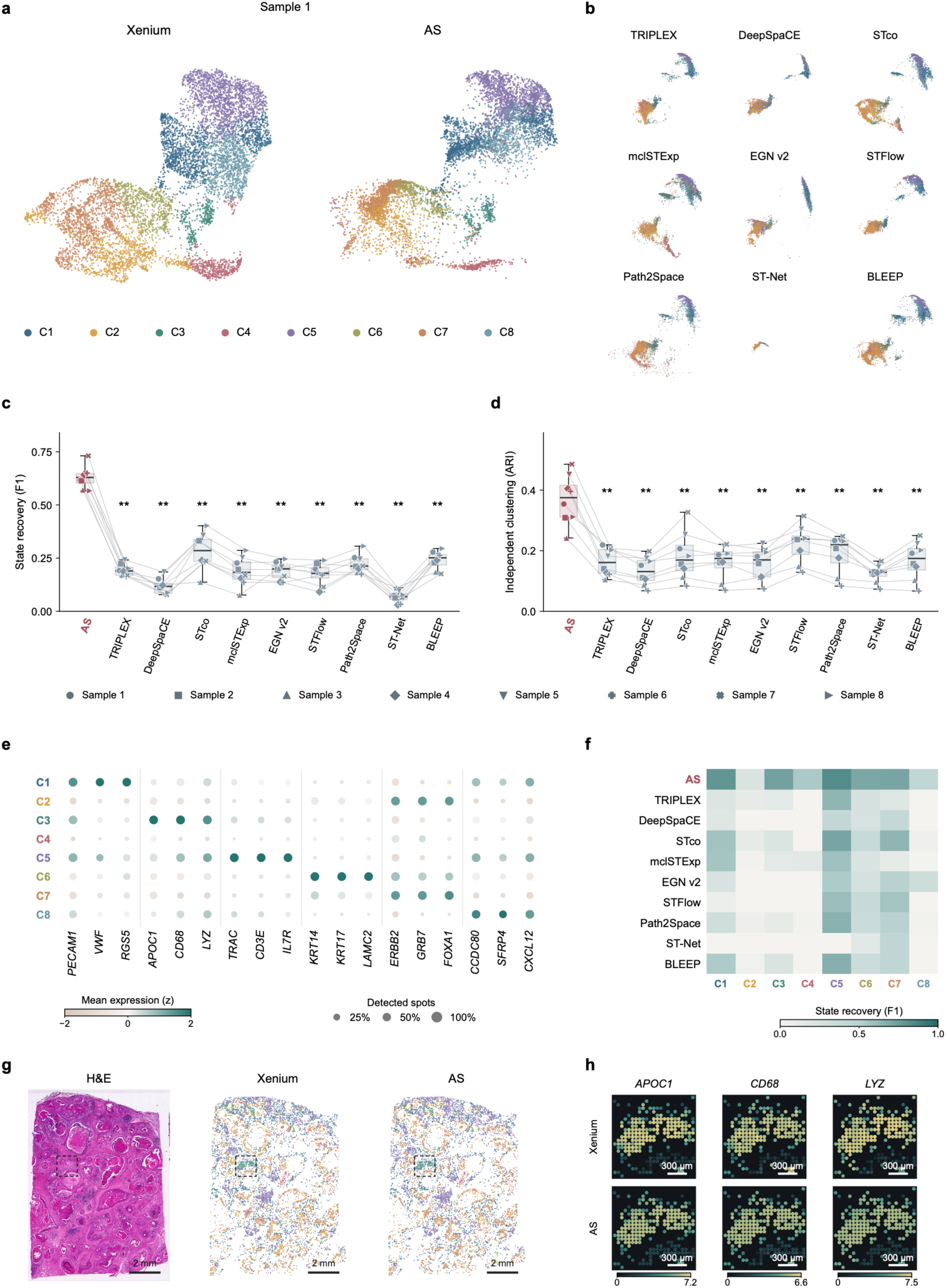
Recovery of molecular states and their spatial organization. **a,b**, Xenium, AS and nine comparator expression profiles from Sample 1 projected into a measured-expression UMAP reference. Colours indicate eight measured reference states and remain fixed for corresponding spots across methods. **c**, Macro-F1 for nearest-centroid recovery of reference states. **d**, Adjusted Rand index (ARI) between independently obtained measured and predicted Leiden partitions. In **c,d**, points represent eight patients; symbols and lines identify paired observations. Boxes show medians and interquartile ranges; whiskers indicate minima and maxima. Two-sided exact paired Wilcoxon tests compare each method with AS. Benjamini–Hochberg correction covers 27 comparisons across F1, ARI and high-dimensional neighbourhood overlap; ** denotes 0.001 ≤ *q* < 0.01. **e**, Measured marker profiles of Sample 1 states. Colour indicates gene-wise standardized mean expression; dot size indicates the fraction of spots with detected expression. **f**, State-specific recovery F1 in Sample 1. **g**, H&E and measured or predicted state distributions; scale bars, 2 mm. **h**, Local measured and predicted expression of *APOC1*, *CD68* and *LYZ*, with shared colour limits for each gene; scale bars, 300 µm. The region was selected using measured C3 density. Analyses use matched 55-µm spots; AS denotes multi-tissue AS. Reference projections are descriptive, whereas ARI is calculated independently of UMAP.

This visual correspondence was supported by quantitative analyses that did not depend on the two-dimensional layout. AS achieved a mean macro-F1 of 0.630 for reference-state recovery, compared with 0.283 for the strongest comparator, STco (Fig. 5c). We also constructed separate neighbourhood graphs and performed independent Leiden clustering on measured and predicted expression^42^. AS achieved a mean adjusted Rand index of 0.369, compared with a maximum of 0.223 among external methods (Fig. 5d). Both advantages were consistent across all eight patients (Benjamini–Hochberg-adjusted q = 0.0078 for every comparison). Thus, the improvement extended beyond individual-gene agreement to the recovery of multigene expression structure.

The recovered states also carried interpretable biological information. In Sample 1, measured profiles distinguished vascular-associated expression of *PECAM1* and *VWF*, myeloid-associated *APOC1*, *CD68* and LYZ, T-cell-associated *TRAC, CD3E* and *IL7R*, and distinct epithelial programmes (Fig. 5e). These patterns were consistent with the cellular diversity described in breast cancer atlases^43^. AS recovered the myeloid-associated C3 and T-cell-associated C5 states with F1 scores of 0.727 and 0.829, respectively, although recovery remained uneven across states (Fig. 5f). Mapping these assignments back to tissue reproduced broad spatial domains, while local expression maps retained the distribution of the C3-associated genes *APOC1*, *CD68* and *LYZ* (Fig. 5g,h). Together, these results support the recovery of biologically interpretable molecular states and their tissue distribution, rather than correlation gains alone.

### Alpha Spatial recovers the spatial and molecular features of *POSTN*-associated fibroblast niches

Having established the recovery of molecular states, we next asked whether Alpha Spatial could reconstruct a biologically coherent stromal niche. Fibroblasts adopt distinct transcriptional states across healthy and diseased tissues^44^. In breast cancer, Null et al. showed that periostin-expressing stromal cells organize collagen and support collective invasion and lymphatic dissemination^45^. This provided a biological framework for testing whether histology-derived predictions recovered both a *POSTN*-associated molecular programme and its spatial relationship to epithelial cells. We examined two breast cancer sections with paired Xenium measurements, assigning cell identities and *POSTN*-high and *POSTN*-low fibroblast groups separately in measured and predicted expression.

Both data sources placed *POSTN*-high fibroblasts closer to epithelial cells than *POSTN*-low fibroblasts (Fig. 6a,b,d). Depth-stratified regional contrasts were negative in both sections for Xenium and AS (all adjusted q < 0.001). This spatial association was accompanied by higher expression of *LRRC15, MMP2, CTHRC1, INHBA* and *MMP11* in the *POSTN*-high group in both measured and predicted profiles (Fig. 6c,e). Thus, the recovered distinction extended beyond POSTN itself to a coordinated set of stromal-state and remodelling-associated genes.

**Fig. 6.**
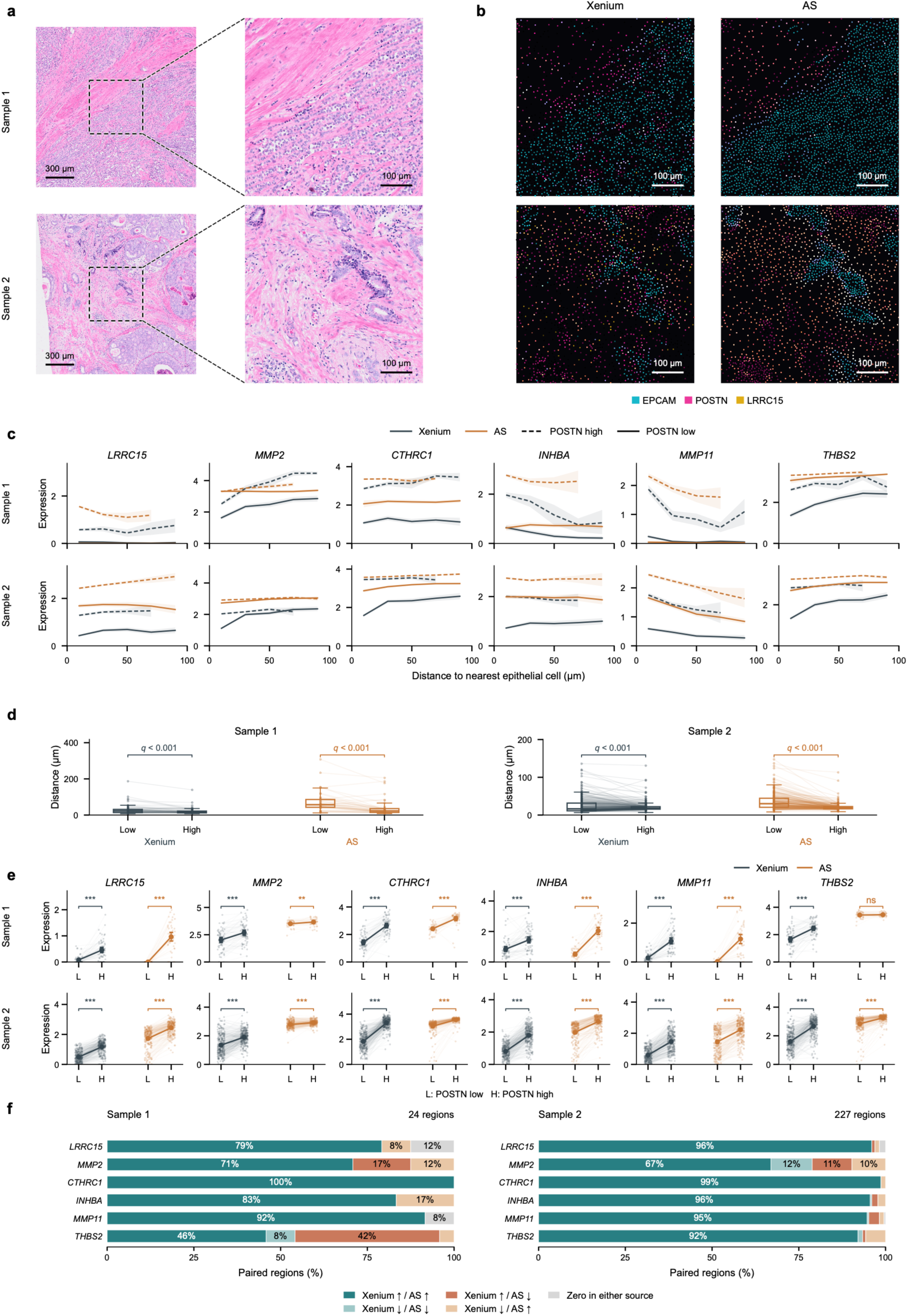
Recovery of POSTN-associated stromal organization. **a**, H&E views and corresponding enlargements from two breast cancer sections; scale bars, 300 µm and 100 µm. **b**, Matched Xenium and AS pseudocolour expression maps of *EPCAM*, *POSTN* and *LRRC15*, using shared gene-specific scales within each section; scale bars, 100 µm. **c**, Fibroblast expression versus distance to the nearest epithelial or basal epithelial cell. Dashed and solid lines denote *POSTN*-high and *POSTN*-low groups; lines show regional means and shading indicates 95% block-bootstrap intervals. **d**, Regional high–low comparisons of epithelial distance. Boxes show medians and interquartile ranges, with 1.5-IQR whiskers. **e**, Regional expression comparisons; large points and error bars show means and 95% block-bootstrap intervals. Connected observations in **d,e** belong to the same 500-µm region; Xenium/AS region counts are 63/43 for Sample 1 and 279/249 for Sample 2. **f**, Joint directions of high–low expression differences across 24 and 227 regions evaluable in both sources; zero denotes a near-zero contrast in either source. Cell identities and *POSTN* groups were defined separately by source: high, above the 75th percentile; low, at or below the 25th percentile. Expression is log1p(CP1,000) over the shared supported gene set. Tests use depth-stratified regional contrasts and 9,999 two-sided sign flips, with Benjamini–Hochberg correction across samples, endpoints, identity definitions and block sizes within each source. *q* < 0.05 \**, < 0.01 **, < 0.001 \*\*\**; NS, not significant. Regions are within-section observations.

The molecular differences were also reproduced locally. Across 24 and 227 matched spatial regions in Samples 1 and 2, respectively, *CTHRC1* increased in the *POSTN*-high group in both sources in 100% and 98.7% of regions; corresponding proportions for *MMP11* were 91.7% and 94.7% (Fig. 6f). Recovery was nevertheless gene-dependent: the *THBS2* contrast was not significant in AS for Sample 1 (q = 0.550), and predicted high–low distance contrasts were larger than measured contrasts (−33.1 versus −8.1 µm and −15.2 versus −3.5 µm). These within-section comparisons show that AS recovered key molecular and positional features of a *POSTN*-associated stromal niche, although the magnitude of spatial differences was not precisely calibrated.

### Alpha Spatial reveals HER2-associated molecular and spatial organization

Before extending Alpha Spatial to clinical cohorts, we examined the spatial scale for analysing coordinated molecular variation. Across eight patients, aggregating cell-level predictions into 55-µm bins improved agreement with measured gene–gene correlations and multivariate expression structure (Supplementary Fig. 5). Spatial aggregation also outperformed occupancy-matched random grouping, supporting the contribution of local tissue organization rather than averaging alone. These results motivated regional aggregation for subsequent molecular-programme and spatial analyses, while retaining cell-level predictions for local expression mapping.

Spatial transcriptomic and multiplexed imaging studies have shown that breast tumour heterogeneity involves both molecular identity and the organization of immune and proliferative compartments^46,47^. We therefore asked whether AS could extend molecular reconstruction from paired experimental specimens to a histological cohort with existing HER2 annotations. Among 185 evaluable cases, predicted *ERBB2* expression was higher in the 90 HER2-positive cases than in the 95 HER2-negative cases (group means, 2.75 versus 2.33; two-sided Welch’s p = 0.0299; Fig. 7a). These case-level estimates averaged absolute predictions within occupied 55-µm bins and then equally across bins, without total-expression normalization. The agreement with HER2 status supported the biological relevance of the predicted molecular signal at the cohort level.

**Fig. 7.**
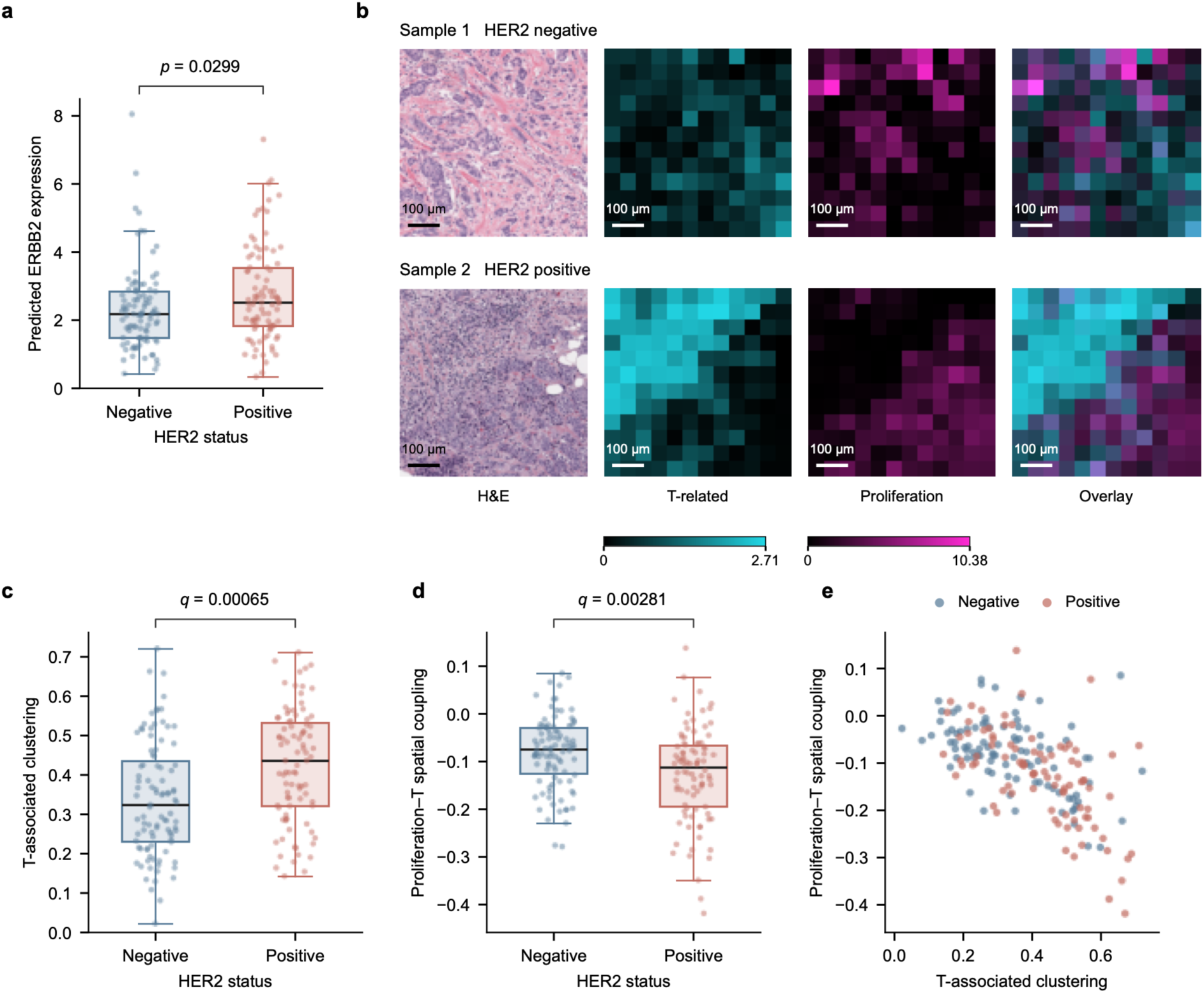
HER2-associated molecular and spatial organization. **a**, Case-level predicted *ERBB2* expression, averaged within occupied 55-µm bins and then equally across bins. **b**, Selected illustrative regions showing H&E, predicted T-associated and proliferation-associated expression, and their overlay. Examples were selected to illustrate spatial contrast; channel scales are shared between cases. Scale bars, 100 µm. T-associated genes: *CD3E*, *TRAC*, *CD8A* and *IL7R*; proliferation-associated genes: *MKI67*, *TOP2A*, *BIRC5*, *CCNB1*, *CDC20* and *UBE2C*. **c,d**, T-associated clustering and proliferation–T spatial coupling, calculated from standardized adjacent-bin products after subtracting within-ROI-stratum random-arrangement expectations. Negative coupling indicates spatial opposition of predicted programmes. **e**, Joint distribution of the two spatial metrics. Quantitative panels include 95 HER2-negative and 90 HER2-positive cases; each point represents one case. Boxes show medians and interquartile ranges; whiskers extend to 1.5 interquartile ranges. Two-sided Welch tests; nominal *p* in **a** and BH-adjusted *q* across 28 spatial comparisons in **c,d**.

We next examined whether these molecular differences were accompanied by differences in spatial organization. Using fixed T-associated and proliferation-associated gene sets, we quantified local clustering and cross-programme coupling on neighbouring 55-µm bins. HER2-positive cases showed stronger T-associated clustering than HER2-negative cases (mean score, 0.423 versus 0.337; Benjamini– Hochberg-adjusted q = 0.000646; Fig. 7c). Proliferation–T coupling was also more negative (−0.128 versus −0.082; q = 0.00281; Fig. 7d), indicating a stronger tendency for the two predicted programmes to occupy spatially opposing neighbourhoods relative to the within-region random-arrangement baseline. Selected histological regions illustrate these contrasting arrangements, with T-associated and proliferative signals occupying distinct local territories in the HER2-positive example (Fig. 7b).

Thus, the cohort-level distinction extended beyond *ERBB2* abundance to the organization of predicted immune-associated and proliferative programmes. The joint spatial profiles nevertheless overlapped between HER2 groups, revealing substantial within-group heterogeneity rather than a binary spatial phenotype (Fig. 7e). These findings support histology-derived molecular maps as a means of studying tumour organization across clinical collections.

### Histology-derived spatial states reveal associations with trastuzumab response

We next asked whether molecular maps reconstructed from routine histology could connect local tissue heterogeneity to patient-level treatment outcomes. We analysed 81 evaluable patients from the Yale trastuzumab-response cohort, comprising 33 responders and 48 non-responders^48^. From 418,378 spatial bins at 55-µm resolution, we identified an epithelial-associated compartment and reclustered its 211,270 bins after Harmony integration^49^. Four molecular states captured matrix-associated, *CXCL14*-associated, epithelial-high and low-expression mixed profiles (Fig. 8a–c). Because these bins contain multiple cells, we interpreted the states as local molecular environments rather than pure epithelial subtypes.

**Fig. 8.**
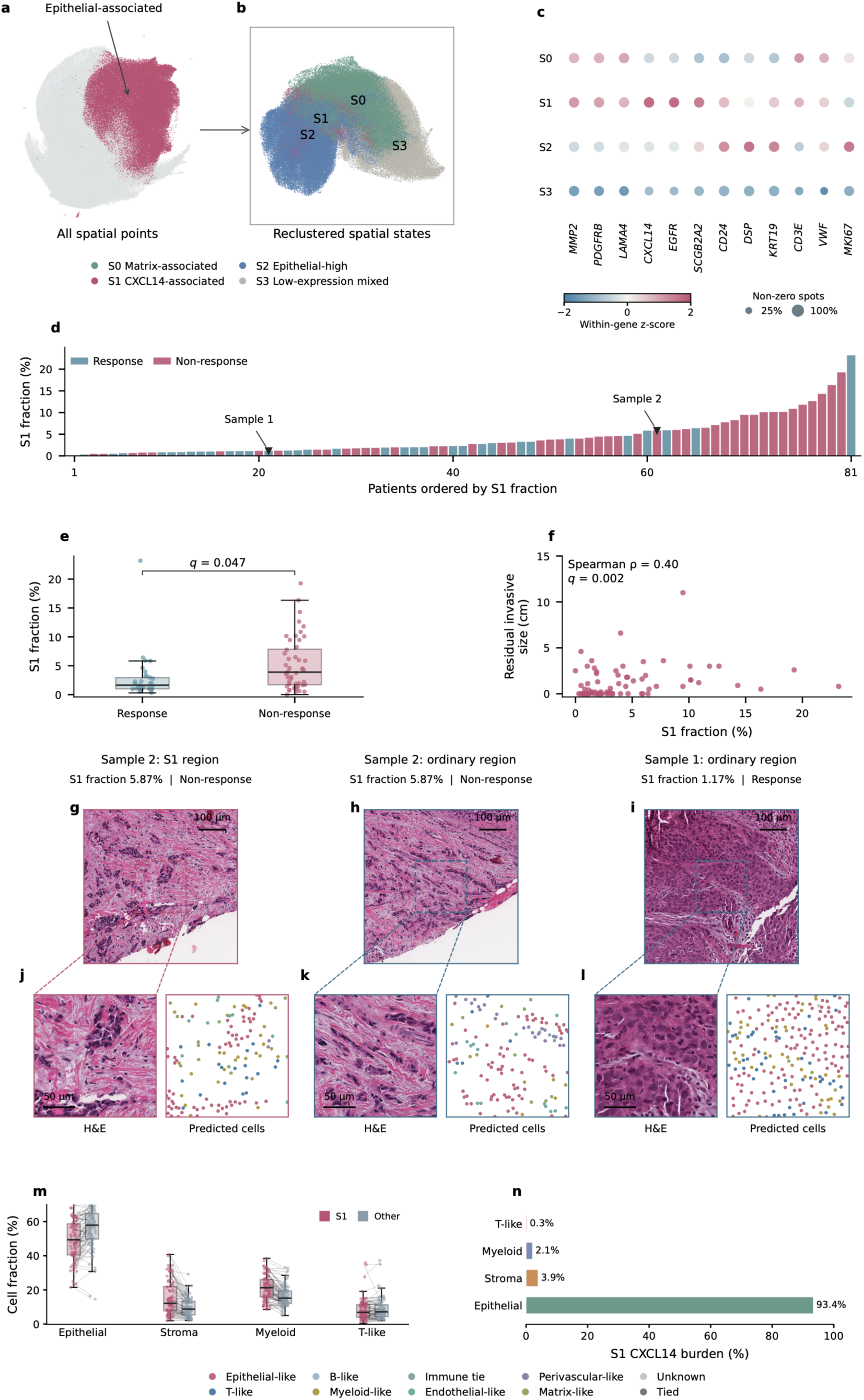
Histology-derived spatial states and trastuzumab response. **a,b,** Global UMAP of 418,378 55-µm spatial bins (a) and reclustering of 211,270 epithelial-associated bins into four states (b). **c,** Marker profiles; colour denotes within-gene standardized patient-mean log1p expression, and dot size denotes the non-zero fraction. **d,** Patient-ranked S1 abundance, defined relative to all epithelial-associated bins. **e,f,** S1 abundance by response (33 responders, 48 non-responders; e) and versus residual invasive tumour size (79 patients; f). Tests were two-sided Welch’s t-tests (e) and Spearman correlations with 9,999 permutations (f), with Benjamini–Hochberg correction across four states separately for each endpoint. **g–i,** Histological contexts: an S1 region and an ordinary region from Sample 2, and an ordinary region from Sample 1. **j–l,** Corresponding central zooms with H&E and predicted cell identities; unknown and tied assignments are retained. Ordinary denotes other parent states, not normal tissue. Context and zoom widths are 600 and 200 µm; scale bars, 100 and 50 µm, respectively. In e, points represent patients; boxes show medians and interquartile ranges, with 1.5-interquartile-range whiskers. **m,** Predicted cell-type proportions in S1 and other epithelial-associated regions across 80 patients containing both categories. Points represent patient-level proportions; lines connect measurements from the same patient. Boxes show medians and interquartile ranges, with 1.5-interquartile-range whiskers. Stroma denotes matrix-stromal-like cells. **n,** Contributions of the indicated predicted cell classes to total predicted *CXCL14* expression within S1. Percentages were calculated from summed cell-level predictions pooled across patients, with all cell classes included in the denominator.

The *CXCL14*-associated state, S1, was present in 80 of 81 patients, but its relative abundance varied substantially (Fig. 8d). S1 occupied a mean of 5.25% of epithelial-associated bins in non-responders, compared with 2.78% in responders (difference, 2.47 percentage points; 95% confidence interval, 0.56– 4.38; Benjamini–Hochberg-adjusted q = 0.0475; Fig. 8e). Its abundance also correlated with residual invasive tumour size among 79 patients with evaluable measurements (Spearman ρ = 0.403; q = 0.002; Fig. 8f). Of the four states, only S1 met the 5% false-discovery threshold for either endpoint (Supplementary Fig. 6a,b). The association therefore concerned the extent of this environment within a patient’s tissue, rather than its presence or absence.

We then examined the cellular composition and histological context of S1. Histological views and cell-level molecular maps compared S1-rich regions with other epithelial-associated regions from the same patient and a region from a patient with lower S1 abundance (Fig. 8g–l). Across 80 patients with both regional categories, S1 regions showed higher patient-mean predicted proportions of matrix-like and myeloid-like cells than other epithelial-associated regions (paired differences, +5.13 and +4.93 percentage points), together with a lower epithelial-like proportion (−6.53 percentage points; Supplementary Fig. 6c). These comparisons characterize S1 as a mixed local molecular environment with increased stromal and myeloid representation, rather than a purely epithelial state. Together, the clinical and regional analyses link the burden of this environment to treatment response and residual tumour size.

We next asked whether the cellular composition of S1 differed from that of other epithelial-associated regions and which predicted cell populations contributed most to the *CXCL14* signal. S1 regions contained a lower epithelial-like fraction and higher stromal-, myeloid- and T-like fractions than other regions from the same patients (Fig. 8m). However, when the predicted *CXCL14* burden within S1 was apportioned across these cell classes, the signal was dominated by epithelial-like cells (93.4%), with smaller contributions from stromal-like cells (3.9%), myeloid-like cells (2.1%) and T-like cells (0.3%; Fig. 8n). Thus, S1 represents a mixed local environment in which stromal and myeloid populations are enriched, whereas the reconstructed *CXCL14* signal itself is predominantly epithelial-associated.

## Discussion

Alpha Spatial connects two central challenges in molecular prediction from histology: learning from heterogeneous cell-resolved measurements and constraining the molecular interpretation of morphology. By representing biological context and assay coverage explicitly, AS allows observations collected across tissues and gene panels to contribute to a shared prediction framework. The findings establish complementary benefits of this formulation. Appropriate context improves correspondence with measured expression, while broader tissue experience improves prediction within a single organ. Together, these advances suggest that molecular reconstruction benefits not only from more detailed images or larger training collections, but also from representing the biological and measurement conditions under which image–expression relationships are learned.

Heterogeneous molecular coverage is fundamental to this learning problem. Earlier approaches established expression prediction from paired histology and spatial measurements, as illustrated by ST-Net^13^, while GHIST extended directly supervised prediction to individual cells^50^. More recently, STPath broadened generative learning across organs and technologies^51^. As cellular measurements accumulate, however, their value depends on whether models can retain supervision that is available in some panels but absent from others. Restricting training to intersecting genes discards valid observations, whereas treating unmeasured genes as zeros confounds measurement coverage with biological absence. Integration frameworks such as uniPort demonstrate the value of retaining shared and dataset-specific features^52^, and Nicheformer extends shared representation learning across dissociated and spatial assays^53^. AS brings this principle to image-conditioned quantitative prediction: panel descriptions identify the measurement setting, and named gene–count outputs accommodate different supervision sets without requiring a separate fixed output vector for each panel. Heterogeneous datasets can therefore provide complementary training evidence rather than being reduced to their common denominator.

Language conditioning also provides a way to examine how biological context shapes molecular inference. Visual–language and visual–omics models, including CONCH and OmiCLIP, have established productive connections between histological representations and semantic or molecular descriptions^54,55^. In AS, language specifies the setting in which a quantitative cellular prediction is requested. The prompt experiments show that this setting matters beyond sensitivity to an incorrect label: matched context improved expression correspondence relative to omitted context, whereas incompatible context reduced it. Moreover, the changes involved coordinated contractile, basement-membrane and myeloid-associated signals, indicating that context influenced functionally structured molecular predictions rather than only their overall magnitude. Permuting predicted profiles among cells preserved their distributions and shared context but abolished their correspondence with measured expression. Thus, a patient-level expression background alone could not account for the observed cell-level correlations. These findings support treating histology-to-expression prediction as a biologically ill-posed inverse problem in which cellular image evidence and contextual constraints contribute complementary information.

Cross-tissue learning provides the biological experience from which these conditional relationships can be learned. Multi-organ atlases show that shared cellular identities coexist with tissue-dependent expression^56^, while cross-tissue studies of fibroblasts identify conserved programmes alongside specialized and disease-associated states^44^. Within a restricted training setting, features associated with a particular molecular programme may also accompany other, locally correlated properties. Additional tissues provide examples in which these relationships differ, potentially helping the model distinguish shared cellular attributes from context-dependent associations. The improvement in breast tissue after multi-tissue training is consistent with this interpretation. The transferable representations obtained through broad pathology pretraining, exemplified by UNI, provide a related precedent^32^. Context conditioning and training diversity therefore address different aspects of the same problem: one specifies the biological setting of a prediction, whereas the other expands the experience available to interpret it. The capacity to use heterogeneous supervision is consequently part of the framework’s practical advantage.

The study also distinguishes the resolution of a prediction from the scale at which it supports reliable biological analysis. Accurate prediction of individual genes does not automatically ensure accurate covariance among genes, which is required for multivariate state identification. In the paired scale comparison, local aggregation improved agreement with measured gene–gene relationships and expression representations. Its advantage over occupancy-matched random grouping further supported a contribution from local tissue organization rather than averaging alone (Supplementary Fig. 5). This motivates a multiscale strategy: retain cell-level predictions for local expression mapping, identify coordinated molecular environments at a more stable regional scale, and return to the constituent cells to examine their composition. Such an approach is well suited to the multicellular structures implicated by spatial studies of breast tumour microenvironments^6^. Cell-resolved prediction and regional analysis are therefore complementary, with the analytical scale determined by the biological relationship being examined.

The biological applications illustrate how this strategy connects molecular reconstruction to tissue-level questions. Recovering *POSTN*-associated stromal programmes and their epithelial relationships extends the relevance of predictions beyond isolated markers, in the context of experimental evidence linking periostin-expressing stromal cells to matrix remodelling and invasion^45^. HER2-associated molecular and spatial differences further connect reconstructed maps with the heterogeneity described by spatial profiling of HER2-positive breast tumours^46^. The response analysis advances a different question: how extensively a molecular environment occupies a patient’s tissue, rather than whether it is present at all. The *CXCL14*-associated state was nearly ubiquitous, yet its burden was associated with non-response and residual tumour size. Its mixed epithelial, matrix-like and myeloid-like composition makes the local environment, rather than *CXCL14* alone, the relevant biological unit. This interpretation is informed by the identification of *CXCL14-*expressing matrix-associated fibroblasts in a mouse breast tumour model^57^, and by experimental evidence that stromal organization can restrict trastuzumab-induced immune killing^58^. Recent studies provide complementary support for this interpretation, while also defining its current evidentiary boundary. In HER2-positive breast cancer, SPP1-positive tumour-associated macrophages have been associated with poor response to trastuzumab-based neoadjuvant therapy and shown experimentally to impair NK- and CD8⁺ T-cell function and trastuzumab-dependent cytotoxicity^59^. In a different tumour and treatment context, an integrated single-cell and spatial study of clear-cell renal-cell carcinoma identified a *CXCL14*-containing multicellular programme associated with immune-checkpoint blockade resistance and provided functional evidence for *CXCL14*-associated recruitment of tumour-associated macrophages^60^. These studies support two separate links: myeloid-cell states can influence anti-HER2 treatment response, and *CXCL14*-containing tumour environments can organize immunoregulatory myeloid populations. They do not establish that epithelial-derived *CXCL14* recruits myeloid cells or causes trastuzumab resistance in breast cancer. Together, these convergent but non-identical observations motivate a testable hypothesis that the extent and composition of particular tissue environments contribute to variation in treatment response. In our data, the reconstructed *CXCL14* burden was predominantly assigned to epithelial-like predictions, whereas stromal-like and myeloid-like populations were enriched within S1. This pattern is compatible with an epithelial-associated *CXCL14*– myeloid spatial niche, but spatial co-occurrence does not establish secretion, receptor-mediated recruitment or a causal role in trastuzumab resistance. Orthogonal spatial assays and functional perturbation will be required to test this mechanism.

Several limitations define the scope of these conclusions and the next experiments needed. Detailed paired validation remains concentrated in breast tissue and within the evaluated gene coverage; broader tissues, panels and segmentation-derived targets require additional validation. Gains in quantitative accuracy did not uniformly improve zero/non-zero discrimination, and regional agreement does not establish equivalent accuracy for every constituent cell. Resource-matched studies are needed to separate the contributions of tissue diversity, training volume and domain-knowledge pretraining. Likewise, the prompt experiments establish useful contextual conditioning but do not uniquely identify how semantic knowledge and learned expression priors contribute to it. Evaluating missing, uncertain and incorrect metadata across tissues would clarify the robustness of this interface. Finally, the treatment-associated state requires independent cohorts with frozen definitions and orthogonal molecular measurements to establish reproducibility and clinical value. Perturbational studies would be needed to determine whether the state, or any of its constituent programmes, participates causally in resistance.

AS provides a route for accumulating cell-resolved molecular knowledge across heterogeneous experiments and applying it to routine histology. Its central contribution is to make both the available supervision and the conditions of molecular interpretation explicit. Coupled with analysis at appropriate spatial scales, this approach broadens the use of archived tissue from estimating individual molecular signals to investigating how molecular environments are organized, how they vary between patients and how they relate to treatment outcomes.

## Methods

### Model formulation and architecture

Alpha Spatial (AS) generates cell-associated gene counts from an H&E image patch I, target-cell box b and instruction c specifying tissue or disease, assay panel and task. For a serialized response S of T tokens, the model learns

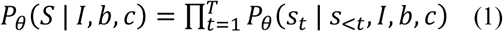

where θ denotes trainable parameters. Expression responses contain non-zero gene–count pairs in original assay units. Panel identity supplies measurement context, not a list of specimen-specific supervision genes; unmeasured genes are not assigned artificial zero targets.

The Qwen3.5-2B backbone^25^ comprises a visual encoder, multimodal projector and autoregressive language model. The visual encoder has 24 layers, hidden width 1,024 and output width 2,048; the language model has 24 layers of width 2,048, with three linear-attention layers followed by one full-attention layer per four-layer group. Projected image features and instruction tokens share the language representation, without a fixed-dimensional gene-regression head. The tokenizer was expanded with 3,740 gene symbols, each verified to have a single-token encoding.

### Training resources and grounded examples

Paired histological and Xenium resources comprised 49 specimens from 48 patients across 29 releases; two lung specimens shared a patient (Supplementary Table 1). The principal model used 40 specimens, excluding nine breast specimens from the top, middle and bottom of Sections 2–4; breast testing included eight independent patients. This checkpoint supported prompt, benchmark, molecular-representation and Yale analyses. Training-resource comparisons additionally used a breast-only checkpoint, also used in both cell-segmentation arms. POSTN analyses used a checkpoint additionally excluding specimens 16_1 and 16_2.

Supervision genes were the union of the top 200 highly variable genes and spatially variable genes with Moran’s I > 0.05 and nominal *p* < 0.05. Selection excluded controls, required detection in at least three cells and at least one count per cell, and used counts scaled to 10,000 followed by log1p transformation; Squidpy spatial statistics used six-neighbour graphs. Targets retained original integer counts. Answers included positive counts only from the selected genes: selected measured zeros were omitted, while unmeasured and measured-but-unselected genes supplied no quantitative target. Prompts named the assay panel without enumerating the selected subset.

Xenium boundary vertices were converted from micrometres using 0.2125 µm/pixel and mapped to H&E by the inverse supplied alignment matrix. Non-overlapping 512 × 512-pixel patches required >60% of pixels below grayscale intensity 215 and at least five complete cell boxes. Boxes were contracted by 20% at each edge, retaining 60% of each dimension, then rounded to patch-relative integer coordinates from 0 to 1,000. Augmentation used hue perturbation of ±5 OpenCV units, independent uniform saturation/value multipliers of 0.9–1.1, and uniformly sampled horizontal, vertical, both-axis or no flipping; boxes underwent the same geometry. JPEG quality was 95. Deterministic patch seeds combined seed 20260728, specimen identity and patch coordinates.

### Domain adaptation and spatial instruction learning

Domain pretraining used PubMedQA, NCBI Gene descriptions, PathVQA and PathMMU^61–63^ before quantitative spatial supervision. Text examples supplied biomedical semantics and visual question– answer examples supplied image-grounded pathology supervision. All model parameters, including the visual encoder and projector, remained trainable; direct preference optimization was not used.

Spatial supervision sampled 20% of eligible cells per patch without replacement, rounded down with a minimum of one. Each sampled cell with positive selected-gene counts supplied a grounded expression example. Independently, cells supplied gene-ranking examples with probability 0.5, using isolated-cell crops resized to a 124-pixel shorter side. Patch-level top-ten gene ranking and tissue/disease identification from the top 50 patch genes were each sampled with probability 0.1. Rankings used decreasing counts and alphabetical tie-breaking. Expression-to-tissue examples supplied neither images nor panel identity. All four tasks minimized response-token cross-entropy,

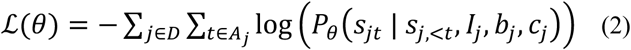

where D is the instruction dataset, *A_j_* comprises response-token positions and *s_jt_* is the target token. Instruction and padding tokens were excluded; no separate count-regression loss was used.

Both stages used three epochs, AdamW (learning rate 1 × 10⁻⁵; β₁ = 0.9, β₂ = 0.999, ε = 10⁻⁸; zero weight decay), cosine decay, 50 warm-up steps, gradient-norm clipping at one, sequence length 4,096, per-device batch size one, bfloat16, sequence packing and seed 42. Domain pretraining used two-step gradient accumulation and ZeRO stage 3; spatial supervision used four-step accumulation and ZeRO stage 2.

### Decoding and matched molecular evaluation

Paired comparisons fixed checkpoint, images and target identities except for the tested factor. Decoding used temperature zero. Gene–count dictionaries were mapped to the measured benchmark panel, POSTN supported-gene intersection or Yale output-feature set. Missing keys within a valid response became zeros only within the specified analysis set; malformed responses, invalid numerical values and length-truncated responses remained missing. Genes outside that set were not appended as zero features.

For gene g in patient p, let *x_ig_* and *y_ig_* denote measured and predicted expression at matched observations i. Gene-wise PCC and log1p RMSE were

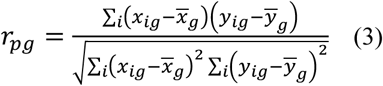

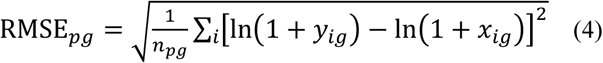

where bars denote observation means and *n_pg_* is the matched count. Patient summaries averaged genes equally and cohort summaries averaged patients equally. Constant-vector PCCs remained undefined.

Primary summaries used each arm’s finite correlations; common-gene analyses and gene-wise paired tests required finite values in both arms. For patient-level error E, relative reduction was

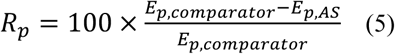

and was averaged across patients; tests used paired error differences. Conditional RMSE used measured-positive cells, background was mean log1p prediction in measured-zero cells, and missed prediction was the fraction of measured-positive cells predicted as zero, each summarized by gene then patient.

To assess spatial allocation, measured and predicted counts were normalized to unit masses 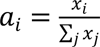 and 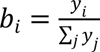 at shared patch-normalized coordinates. For directions *θ_k_* = kπ/32, k = 0,…,31, sorted projections *z_j_* gave

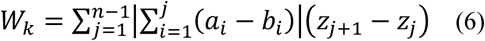

with masses reordered identically. Discrepancy was averaged across directions, eligible patches per gene, then genes per patient. Patches required ≥20 valid cells and positive mass in both modalities. Unit-mass normalization applied only to this metric.

### Context and training-resource comparisons

Three-context analysis used 79,912 common-valid cells from eight patients, substituting Breast cancer, Breast or Lung cancer while omitting panel identifiers. Comparator c versus Breast cancer reference r defined expression shifts and switch rates as

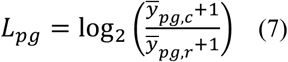

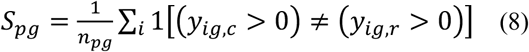

where bars denote cell means and 1[·] is an indicator. Detection shifts were comparator-minus-reference positive-prediction fractions; their associations with reference-minus-comparator PCC were evaluated across genes by within-patient Spearman correlation. A fourth condition omitted tissue, disease and panel information, retaining images, boxes, prediction instruction and decoding settings. Of 80,000 targets, 79,928 yielded valid outputs, 72 invalid JSON and none truncation. Four-condition analysis used 79,869 common-valid cells and, for PCC, genes finite in all four arms. Entire predicted vectors were permuted among cells within patients 1,000 times, sharing each permutation across genes and conditions (seed 20261004 + patient index), preserving distributions while disrupting correspondence.

Prompt-compartment annotations used measured counts scaled to 10,000, log1p-transformed and gene-wise standardized. Mean marker scores assigned the highest-scoring subtype, then its broad lineage; cells without positive annotation markers remained unassigned. Epithelial markers were EPCAM/KRT19/ESR1 and KRT5/KRT14/KRT17; stromal markers were ACTA2/TP63/KRT5, FBLN1/SFRP4/SFRP1, ACTA2/TAGLN/CNN1 and PECAM1/VWF/FLT1; immune markers were CD3E/CD8A/TRAC, MS4A1/BANK1/TNFRSF13C, CD68/CD163/LYZ and KIT/CPA3. Evaluation excluded annotation genes and required ≥30 cells per compartment, without spatial smoothing. Displayed expression was log1p-scaled by each gene’s maximum across measured and all contextual predictions in the displayed field; normalized error reduction was comparator absolute error minus Breast cancer absolute error on that scale.

Training-resource comparisons used 79,980 matched cells from eight patients, fixed functional modules and source cell annotations. Within-type tests required ≥100 cells per patient/type, ≥3 finite module-gene PCCs in both arms and ≥4 paired patients; cell thresholds of 50 and 200 were sensitivity analyses. Auxiliary support counted distinct non-breast tissues selecting each gene. Patient-specific regressions related PCC gain to standardized support breadth, measured log1p mean, detection fraction and baseline PCC.

### Cross-method benchmarking and cell localization

Comparators were TRIPLEX, DeepSpaCE, STco, mclSTExp, EGN v2, STFlow, Path2Space, ST-Net and BLEEP, using official components with coverage-aware adaptations and method-specific schedules supplied with the code. Comparator supervision used designated breast resources; multi-tissue AS additionally used heterogeneous resources. The benchmark compared trained frameworks, not architecture under identical training exposure.

Common support comprised 280 genes and 26,068 mutually evaluable circular 55-µm capture regions across eight patients. AS counts were summed within regions. Each measured or predicted vector v used its own panel total for transformation,

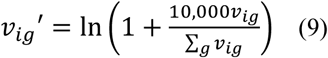

without rescaling predictions to measured depth. Pooled measured–predicted regressions included an intercept; R² = 1 − SSE/SST used predicted values as the dependent variable. Spatial-variation ratios were gene-wise SD(predicted)/SD(measured), summarized by the within-patient median and equal-patient mean. Expression and detection quintiles were defined from reference data per patient.

Cellpose-SAM supplied image-derived masks in one breast section; the inference interface accepts other segmentation methods. Mutual maximum-IoU matching within shared 512-pixel patches used geometry, not expression. IoU ≥0.30 and centroid separation ≤10 µm retained 6,191 of 10,000 reference cells and 279 genes; IoU thresholds 0.10 and 0.50 retained 7,526 and 2,181 matches. Both arms used the breast-only checkpoint and 2,048-token limit. Uncertainty used 1,000 paired resamples of 53 spatial blocks (1,024 × 1,024 pixels; seed 20260923). Box construction and image encoding differed, so this was not an isolated coordinate perturbation.

### Molecular representations and spatial scale

Within each benchmark patient, reference-fitted gene standardization, PCA (30 components) and UMAP (30 neighbours, minimum distance 0.3, Euclidean distance) defined a common projection^41^. Predictions were assigned to the nearest of eight reference k-means centroids in PCA space. Macro-F1 was

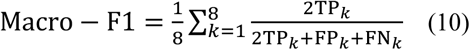

where TP, FP and FN are state-specific true-positive, false-positive and false-negative counts. Independent symmetrized, unweighted 30-neighbour graphs underwent Leiden RB clustering (resolution 1, two iterations, seed 20260927)^42^. For partition contingency counts *n_ab_*, row totals *r_a_*, column totals *c_b_* and N observations, set 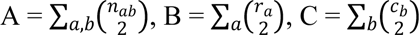 and 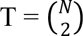; adjusted Rand index was

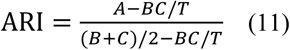

Neighbour overlap was

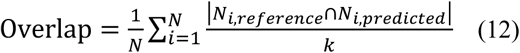

for neighbour sets N_*i*_ in standardized gene space, with k = 30 and sensitivities 15 and 50. States were patient-specific; metrics were calculated once per patient.

Scale analysis compared multi-tissue predictions with Xenium counts for 1,733,702 identity-matched cells from eight patients. Square grids of 27.5, 55 and 110 µm originated at (0,0), with a 55-µm grid additionally shifted 27.5 µm along both axes. Occupied bins averaged absolute measured and predicted counts separately; the unshifted 55-µm grid contained 114,686 bins. All valid cells were retained. Genes required log1p SD >10⁻¹⁰ in both modalities at every scale/origin, retaining 280 genes except Sample 5 (279). No total-expression normalization or Harmony correction was applied.

After log1p transformation of bin means, gene–gene correlation error compared measured and predicted matrices R and R^ over G genes:

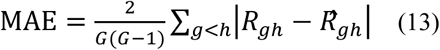

For multivariate agreement, matrices were independently gene-centred, divided by population SDs and reduced independently to 30 PCs. Centred score matrices U and V yielded linear centred kernel alignment,

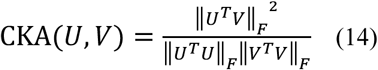

where F denotes the Frobenius norm. State assessment instead fitted standardization, 30-PC PCA and eight MiniBatchKMeans states to measured data and applied them unchanged to predictions (seed 20261004, ten initializations, batch size 4,096, maximum 100 iterations). Macro-F1 used scale-specific states. Variance retention was the median gene-wise ratio of bin-mean to cell-level population variance in absolute counts.

Occupancy-matched controls shuffled measured–predicted cell pairs jointly within patients before grouping, preserving pairing and exact bin sizes. Ten randomizations per scale used seeds 20261004 + 1,000 × patient index (1–8) + replicate index (0–9); metrics were averaged within patients. Correlation error and CKA at 55 µm versus cells formed a two-test Holm family; spatial versus random comparisons of these metrics formed another. Other scale metrics were descriptive. Mean differences used 20,000 paired patient bootstraps (seed 20261004).

### POSTN-associated stromal organization

Sections 16_2 and 16_1 contained 365,601 and 574,527 paired-valid cells and 272 and 310 supported genes. Both modalities used each section’s supported-gene universe. Cells required ≥20 supported counts; expression was scaled to 1,000 and log1p-transformed. Marker values were divided by the positive-reference-cell 95th percentile and clipped to [0,1]. Fixed module means assigned the unique highest positive identity; zeros/ties remained unresolved. Predictions reused reference scales but received independent identities. POSTN and remodelling endpoints were excluded from annotation.

Within each modality/section, fibroblasts above the *POSTN* 75th percentile and at/below the 25th percentile formed high and low groups; intermediate cells were retained outside this contrast. Endpoints were *LRRC15, MMP2, CTHRC1, INHBA, MMP11, THBS2* and distance to the nearest epithelial/basal-epithelial centroid. Expression-depth quartiles within 500-µm blocks required ≥5 high and ≥5 low cells; contrasts were weighted by the smaller group size within blocks, then equally across blocks. Tests required ≥8 blocks and used 9,999 sign flips and 2,000 block bootstraps. Sensitivities used 1,000-µm blocks and reduced-marker identities. Distance curves used 20-µm bins, ≥5 cells per block/bin, ≥5 blocks and 1,000 block bootstraps.

### Yale cohorts, prediction and aggregation

HER2-TUMOR-ROIS includes 192 HER2-status cases and 85 trastuzumab-response cases^48^. Status labels used supplied IHC/FISH annotations. Response biopsies preceded neoadjuvant trastuzumab with/without pertuzumab and surgery. Supplied pathological complete response required absence of residual invasive carcinoma, lymphovascular invasion and metastatic carcinoma; residual in situ disease alone did not indicate nonresponse. Residual invasive tumour size was measured in centimetres, not treated as survival. Source-study approval was Yale Human Investigation Committee protocol 9505008219; this study reused public data without recruitment or treatment assignment.

Five status cases lacked ROIs and four response cases had invalid/out-of-image ROIs, leaving 187 and 81 processable cases. Quality-controlled status analysis included 185 cases (90 positive, 95 negative; 7,269,888 valid first-pass cells); response analysis used final resolved outputs from 81 patients (33 responders, 48 nonresponders; 5,866,733 valid cells).

NuLite-H segmented supplied ROIs at 0.27381295745691947 µm/pixel using 512-pixel tiles, 384-pixel cores and 64-pixel halos. Exclusive centroid ownership prevented duplicate assignments. ROI unions subtracted negative regions within the same annotation. RGBA images were composited onto white and bilinearly resized. Native coordinates and ROI membership were retained; overlapping identities were deduplicated. Size/stain flags were retained without excluding flagged nuclei.

Both cohorts used the principal checkpoint and identical text:

> Focus on the <|object_ref_start|>cell<|object_ref_end|><|box_start|>[x1,y1,x2,y2]<|box_end|> in this Breast Cancer section. Using the Biomarkers and Housekeeping Genes Custom Panel, predict the cell’s transcriptomic profile. Return non-zero gene counts in descending order as a JSON object.

Inclusive mask-box coordinates were integers on the pixel/512 × 1,000 scale, serialized without spaces. RGB patch and text formed one user message in the checkpoint’s chat template with generation prompt appended. Only image and box varied; case identity and clinical labels were excluded. Decoding used temperature zero, seed 20260927, context length 8,192, output limit 2,048, ≤512 concurrent sequences, ≤16,384 batched tokens and no prefix caching. Stopped malformed responses could receive one identical-input retry; resolved outputs replaced first-pass values only with valid retries. Truncated/unresolved responses remained missing. Status analysis used quality-controlled first-pass outputs; the response atlas used resolved outputs. Valid cells were not filtered by predicted expression, lineage or outcome.

Occupied 55-µm square bins averaged absolute predictions without total-expression normalization, using the slide’s physical origin. Status keys retained exact ROI membership; response bins pooled deduplicated cells at each position within the ROI union. Empty bins were omitted and case means weighted bins equally. Local annotation used mean log1p marker counts with ≥2 positive markers: epithelial score was max(core, basal), immune score max(T, B, myeloid), alongside endothelial, perivascular and matrix scores. The highest supported score assigned identity; unsupported/tied cells remained included.

### HER2-associated spatial organization

ERBB2 equal-bin means were compared across 90 positive and 95 negative cases. Eight programs used arithmetic means of absolute gene predictions; six pairs assessed coupling. Undirected edges linked horizontal/vertical adjacent occupied bins within identical ROI-membership strata. Program values were standardized within strata using population SDs. For n bins and edge set E,

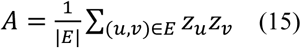

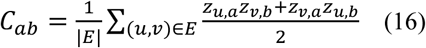

where z denotes standardized expression. Random-row-permutation expectations were −1/(n−1) for A and −mean*i*(*z_ia_z_ib_*)/(n−1) for C. Observed-minus-expected scores were edge-weighted across strata; eligibility required ≥3 bins, an edge and non-constant signals. Average-rank signals and equal-stratum weighting were sensitivities. Welch tests compared groups. Geometry-adjusted models included HER2 status and standardized log bin count, log cells per bin and log(1 + ROI-stratum count), with HC3 covariance. Eight autocorrelations plus six couplings on raw/ranked signals formed a 28-test BH family; geometric adjustment and equal-stratum analyses had separate families.

### Response atlas, state burden and regional composition

The atlas comprised 418,378 bins and 279 features. Outcome-independent selection from 284 positive output keys required detection in ≥1% of valid cells and ≥10 patients; keys began with a letter and contained only letters, digits, underscores, periods or hyphens. Feature filtering did not remove cells. Log1p bin means were gene-centred/scaled by sample SD and reduced to 30 PCs. RAPIDS-singlecell 0.13.5 implemented patient-only original Harmony^49^: θ = 2, ridge penalty 1, σ = 0.1, 100 clusters, block proportion 0.05, maximum 10 outer/200 inner iterations, tolerances 10⁻⁴/10⁻⁵, τ = 0, float64 and seed 20261001. Raw expression remained unchanged. A 15-neighbour NN-descent graph supported Leiden resolution 0.5 and UMAP (minimum distance 0.5, 200 iterations, random initialization); clinical labels and response-associated genes were not embedding features.

Marker-supported epithelial-associated parent clusters supplied 211,270 bins for separate scaling, PCA, Harmony and graph refitting, with Leiden resolution 0.2; the remaining 207,108 bins stayed in the atlas. Markers used patient-paired mean-log1p contrasts against other parent bins and the nearest state by maximum graph connectivity, requiring ≥20 bins in both groups. Candidate markers required positive mean effects and positive effects in ≥70% of evaluable patients. Ranking subtracted each patient’s across-gene mean shift before averaging. Mixed states were not assigned pure identities. Before clinical testing of the final partition, the locally characterized state was matched to an initially inspected candidate by maximum Jaccard overlap; clinical comparisons included all four states.

Patient state burden was

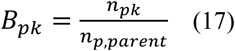

where *n_pk_* and *n_p_*_,*parent*_ count state-k and epithelial-parent bins. Zero burdens were retained and analysed continuously, without outcome-optimized dichotomization. Welch tests compared 33 responders with 48 nonresponders; residual-size Spearman correlations used 79 patients, 9,999 permutations and 2,000 patient bootstraps. Missing endpoints were not imputed.

Cells inherited their bin’s state. Within each patient, predicted-identity percentages used all valid cells in S1 or other epithelial-parent states (S0, S2, S3) as separate denominators, retaining unknown/tied categories. Paired differences were S1 minus other-state percentages. Eighty patients had both categories; the patient without S1 retained zero burden in clinical analyses. Composition summaries were descriptive and patient-weighted. Other-state regions did not denote normal tissue.

### Statistical analysis and reproducibility

Tests were two-sided. Patients were independent biological replicates in breast and Yale comparisons; cells, genes, bins, permutations and within-section blocks did not substitute for patients. The retrospective study used no prospective sample-size calculation or experimental randomization; computational resampling quantified uncertainty, not independent biological repetition.

Prompt tests used paired Wilcoxon signed-rank tests with separate Holm families of three overall, nine lineage, four error/spatial and four shift-association contrasts. In four-condition analysis, Breast cancer versus omitted-context PCC was the single primary, unadjusted test; two other PCC and three RMSE comparisons against omitted context formed a five-test Holm family. These used exact paired Wilcoxon tests and 20,000 patient bootstraps (seed 20261003).

Training-resource tests used exact paired sign tests, excluding zero differences. For n non-zero pairs and k = min(npositive,nnegative),

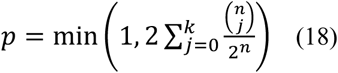

Holm correction covered two primary and four conditional/detection/common-support endpoints separately. Gene-level PCC/error screens used separate BH families; module/within-type families were separate by metric and cell threshold. Benchmark exact paired Wilcoxon tests used BH over 19 comparisons, representation tests over 27, and response burden/residual-size tests over four states separately. POSTN BH families included testable endpoints, identity variants, block sizes and both sections, separately by modality. Holm-adjusted values are labelled *p*, BH-adjusted values *q*.

For group means m, variances s² and sizes n, Welch statistics were

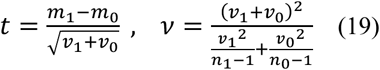

where 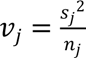. Geometry-adjusted regression used

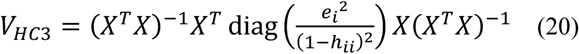

where X is the design matrix, *e_i_* a residual and ℎ*_ii_* leverage, with t inference on n − rank(X) degrees of freedom. Two-sided Monte Carlo tests used

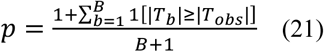

where B is the resample count and *T_obs_*/*T_b_* are observed/resampled statistics. No ANOVA was used.

Patient bootstraps used 20,000 resamples for training-resource, benchmark and representation summaries and 10,000 for prompt-shift correlations; other counts are specified above. Intervals were percentile 95% intervals unless Welch/HC3 intervals were specified. Mean ± s.d. used sample SD. Boxes show median/interquartile range and observed-value whiskers within 1.5 interquartile ranges, except representation boxes (minimum–maximum) and correspondence-permutation boxes (2.5th–97.5th percentiles). Benchmark quintile bands show patient-bootstrap intervals; grey boxes summarize nine external-method means.

### Gene modules and software

Fixed lists below were intersected with supported genes. *POSTN* reduced-marker checks used *CD3E/TRAC* for T cells, *PECAM1/VWF* for endothelial cells and *LUM/PDGFRA* for fibroblasts, excluding POSTN/remodelling endpoints from identity scoring.

### Training-resource comparison modules

| Module | Genes |
| --- | --- |
| Luminal | *ESR1, PGR, GATA3, FOXA1, NAT1, TFF3, SCUBE2* |
| Epithelial | *EPCAM, KRT19, ERBB2, CDH3, CLDN4, KRT5, KRT14, KRT17* |
| Proliferation | *MKI67, TOP2A, BIRC5, CCNB1, CCNE1, CDC20, UBE2C* |
| Lymphoid | *CD3E, CD8A, TRAC, MS4A1, IL7R, TIGIT, CTLA4, CCL5* |
| Myeloid | *CD68, CD163, LYZ, APOC1, ITGAX, S100A8, S100A9* |
| Stromal | *ACTA2, MYH11, CNN1, TAGLN, COL4A1, LAMA2, LAMB1, HSPG2, FBLN1, CXCL12* |

### HER2 spatial programs

| Module | Genes |
| --- | --- |
| HER2-locus pair | *ERBB2, GRB7* |
| Luminal | *ESR1, PGR, FOXA1, GATA3, BCL2* |
| Proliferation | *MKI67, TOP2A, BIRC5, CCNB1, CDC20, UBE2C* |
| Basal | *KRT5, KRT14, KRT17, EGFR, FOXC1* |
| T-cell | *CD3E, TRAC, CD8A, IL7R* |
| B-cell | *MS4A1, BANK1, TNFRSF13C, CD74* |
| Myeloid | *LYZ, CD68, CD163, APOC1* |
| Matrix/stromal | *FBLN1, SFRP4, CCDC80, LAMA2* |

Cross-program pairs were proliferation–T cell, proliferation–myeloid, proliferation–matrix, T cell– myeloid, T cell–matrix and T cell–B cell.

### Yale local cell-annotation modules

| Module | Genes |
| --- | --- |
| Epithelial core | *EPCAM, KRT19, CLDN4, TACSTD2* |
| Epithelial basal | *KRT5, KRT14, KRT17* |
| T | *CD3E, TRAC* |
| B | *MS4A1, BANK1, TNFRSF13C* |
| Myeloid | *LYZ, CD68, CD163* |
| Endothelial | *PECAM1, VWF, PLVAP, CLEC14A* |
| Perivascular | *RGS5, PDGFRB, MYH11* |
| Matrix | *FBLN1, SFRP4, CCDC80, LAMA2* |

### POSTN identity modules

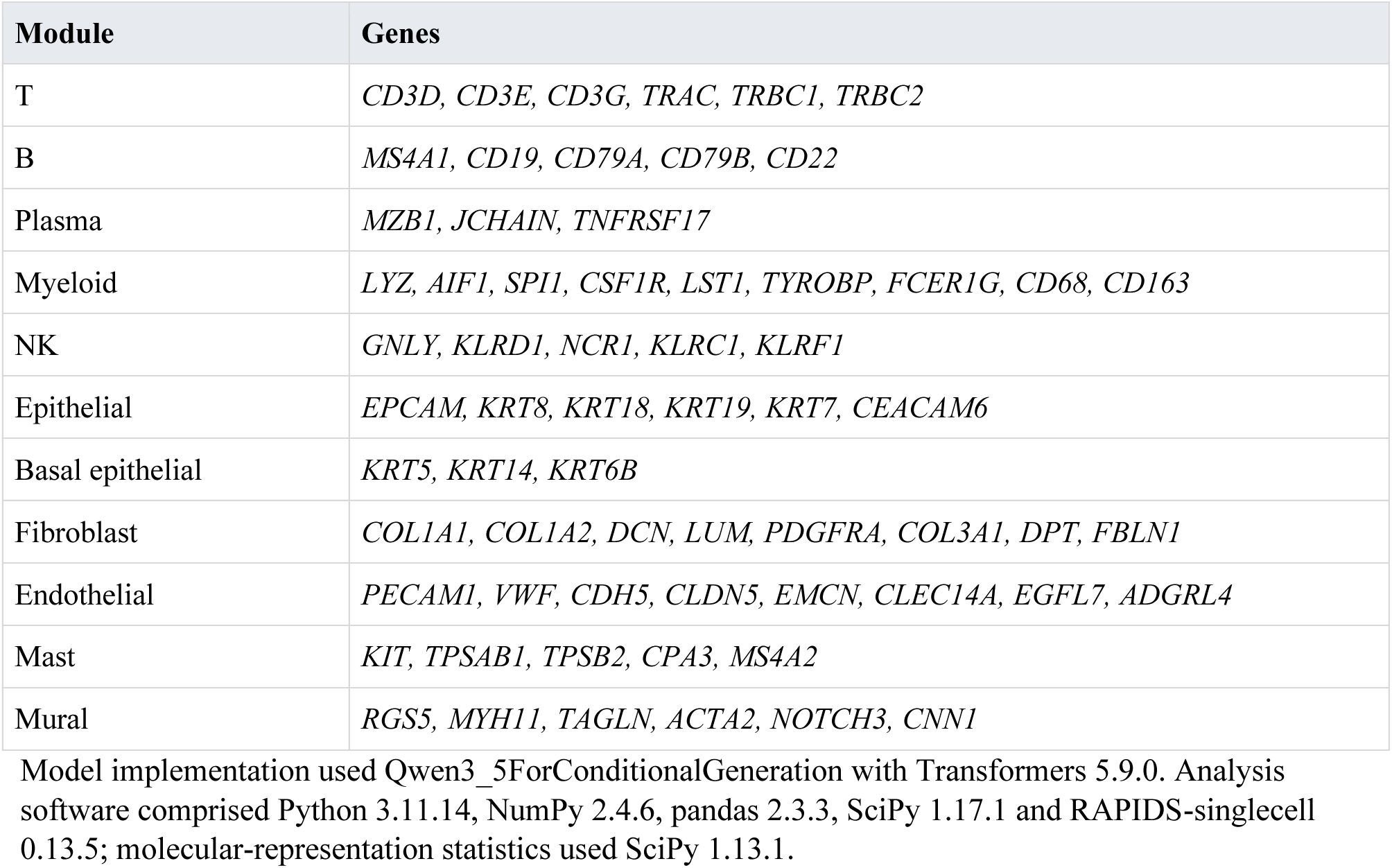

## Data Availability

Publicly available paired histological and Xenium datasets used in this study are listed in Supplementary Table 1, together with their source identifiers and access links. Yale histological images, ROI annotations and clinical labels are available from the HER2-TUMOR-ROIS collection in The Cancer Imaging Archive (https://doi.org/10.7937/E65C-AM96). AlphaSpatial-predicted cell-level expression, spatial coordinates and ROI mappings are deposited in Figshare (https://doi.org/10.6084/m9.figshare.34060014). Deposited expression values are model predictions, not experimentally measured transcript counts. Source data are provided with this paper.

## Code Availability

AlphaSpatial inference code, example inputs, environment specifications and scripts for patient-level treatment-response comparisons are available on https://github.com/xiyangwhu/Alpha-Spatial under the Apache 2.0 licence. The multi-tissue model weights, tokenizer and associated configuration files are deposited in Figshare (https://doi.org/10.6084/m9.figshare.34062837). The repository includes a small inference demonstration and statistical recalculation from released patient-level state counts; it does not currently implement end-to-end reproduction of all manuscript analyses.

## Acknowledgements

We thank Gary Zhang from UCL and Yang Yang, Di Yu from UQ for this study. We thank Hongyu for model development. We thank members of the Liu laboratory for support and discussion. This work is supported by funding from the National Natural Science Foundation of China (NSFC82130051), Tsinghua University-Xiamen Chang Gung Hospital Joint Research Center for Anaphylactic Disease, the Tsinghua-Peking Center for Life Sciences to Y.-C.L. and The CAST Youth Science and Technology Talent Cultivation Program for Doctoral Students.

## Competing declarations

The authors declare no competing interests.

