## Supplementary figure 1-6 for "Alpha Spatial: Language-guided prediction of single-cell spatial gene expression from histology"

### Supplementary Information

**Supplementary Fig. 1 | Context omission and cell-specific prediction correspondence.**

**a,b**, Mean gene-wise PCC (a) and log1p RMSE (b) under four prompt conditions. Context omitted excludes tissue, disease and panel information. Points represent eight independent patients, lines connect paired observations and horizontal marks indicate means. Analyses use 79,869 common-valid cells; PCC uses identical evaluable genes across conditions within each patient. **c**, PCC distributions from 1,000 within-patient permutations of predicted expression profiles against fixed measured cells. Boxes indicate medians and interquartile ranges; whiskers span the 2.5th–97.5th percentiles, with outlying permutations shown individually. **d,e**, Expression shifts versus PCC gains for Breast cancer relative to Breast (d) or Lung cancer (e), using the original 79,912-cell analysis. Points represent patient-mean values for 271 and 272 genes, respectively, each evaluable in at least six patients. Expression shifts are $\log_2$[(comparator mean + 1)/(Breast cancer mean + 1)]; colour indicates comparator-minus-Breast-cancer changes in non-zero cell fractions. Displayed ρ values are means of within-patient Spearman correlations. Tests were two-sided paired Wilcoxon tests. The Breast cancer-versus-omitted PCC comparison was the single primary test; five secondary comparisons used Holm adjustment. Correlation analyses in d,e used a separate four-test Holm family. Source data are provided.

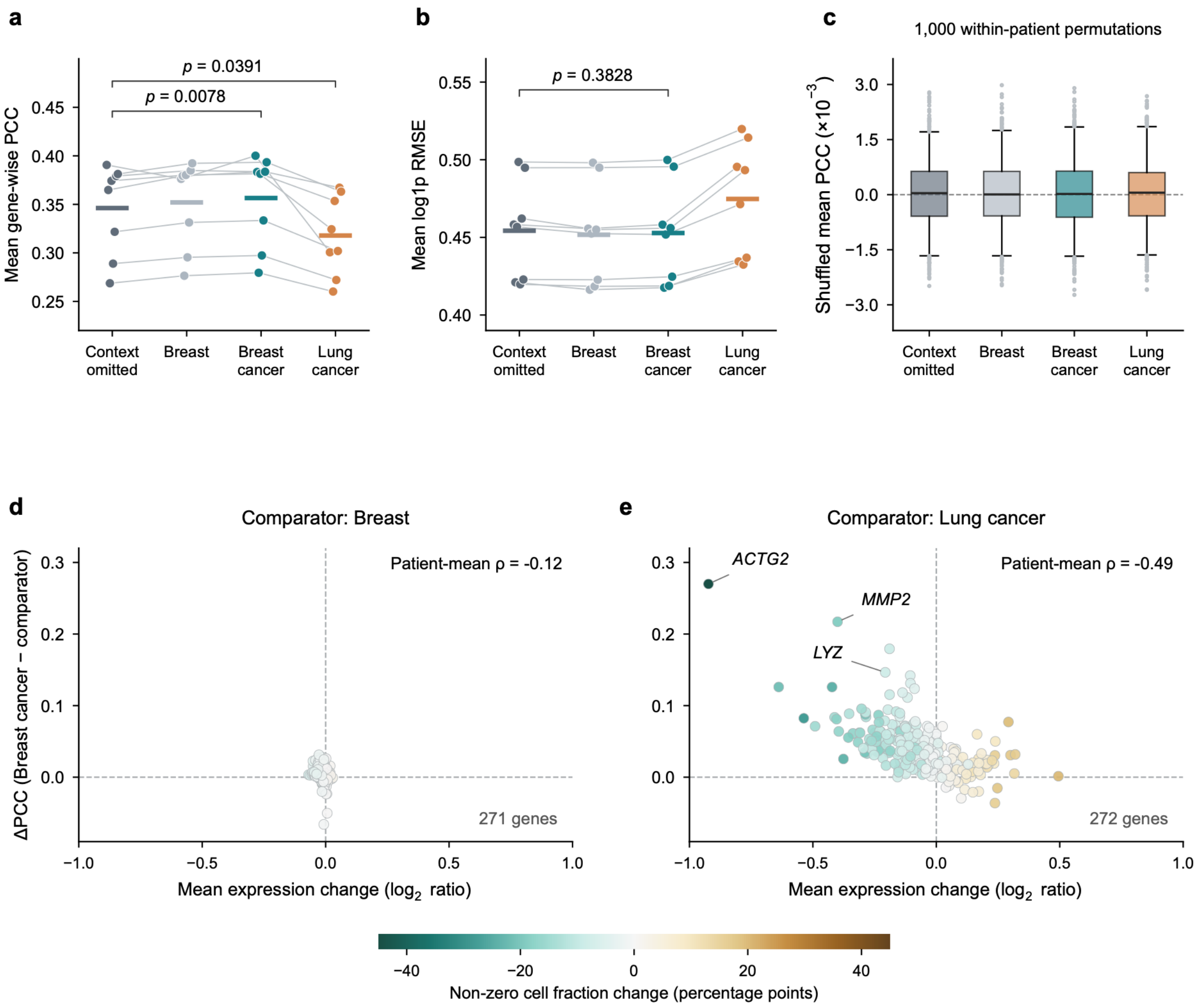


**Supplementary Fig. 2 | Quantitative expression accuracy and detection under multi-tissue learning.**

**a**, Mean log1p prediction in measured-zero cells. **b**, Log1p root mean squared error (RMSE) and missed-prediction rate in measured-positive cells. **c**, Mean gene-wise PCC restricted to genes evaluable in both models (252–280 per patient). Connected points represent eight patients; horizontal bars indicate means. Gene-level metrics are averaged equally within patients. Two-sided exact paired sign tests were Holm-adjusted across four comparisons. **d**, Patient-level PCC differences (multi-tissue minus breast-only) across 280 genes, ordered by average-linkage clustering of their difference profiles; grey denotes undefined correlations.

Analyses use 79,980 matched cells. Measured zero denotes assay non-detection, not established biological absence.

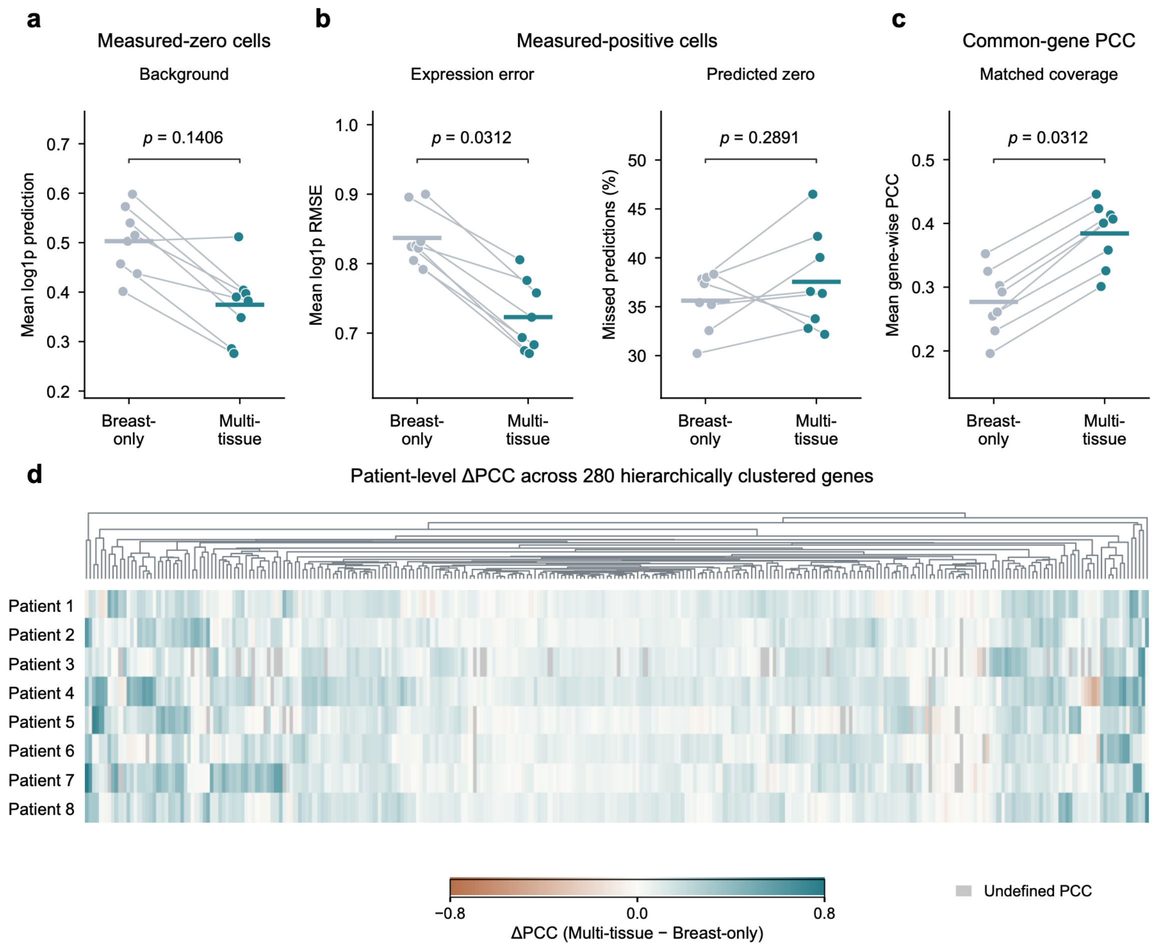


**Supplementary Fig. 3 | Molecular prediction with image-derived cell localization.**

**a**, Two haematoxylin and eosin (H&E) fields with Xenium and image-derived cell segmentation masks; scale bars, 25 µm. **b**, Gene-wise PCC for Xenium-guided and segmentation-guided AS using the breast-only checkpoint (279 genes; 6,191 matched cells from one breast section). Solid and dashed lines indicate the linear fit and identity, respectively; $R^2$ describes agreement between gene-wise PCCs. **c**, Matching coverage among 10,000 reference cells and cumulative centroid-distance and intersection-over-union (IoU) distributions for 7,621 mutual-best-overlap pairs. Accepted matches require IoU $\geq$ 0.30 and centroid distance ≤ 10 µm. Matching uses geometry, not expression. Mask agreement is not segmentation accuracy against manual annotations.

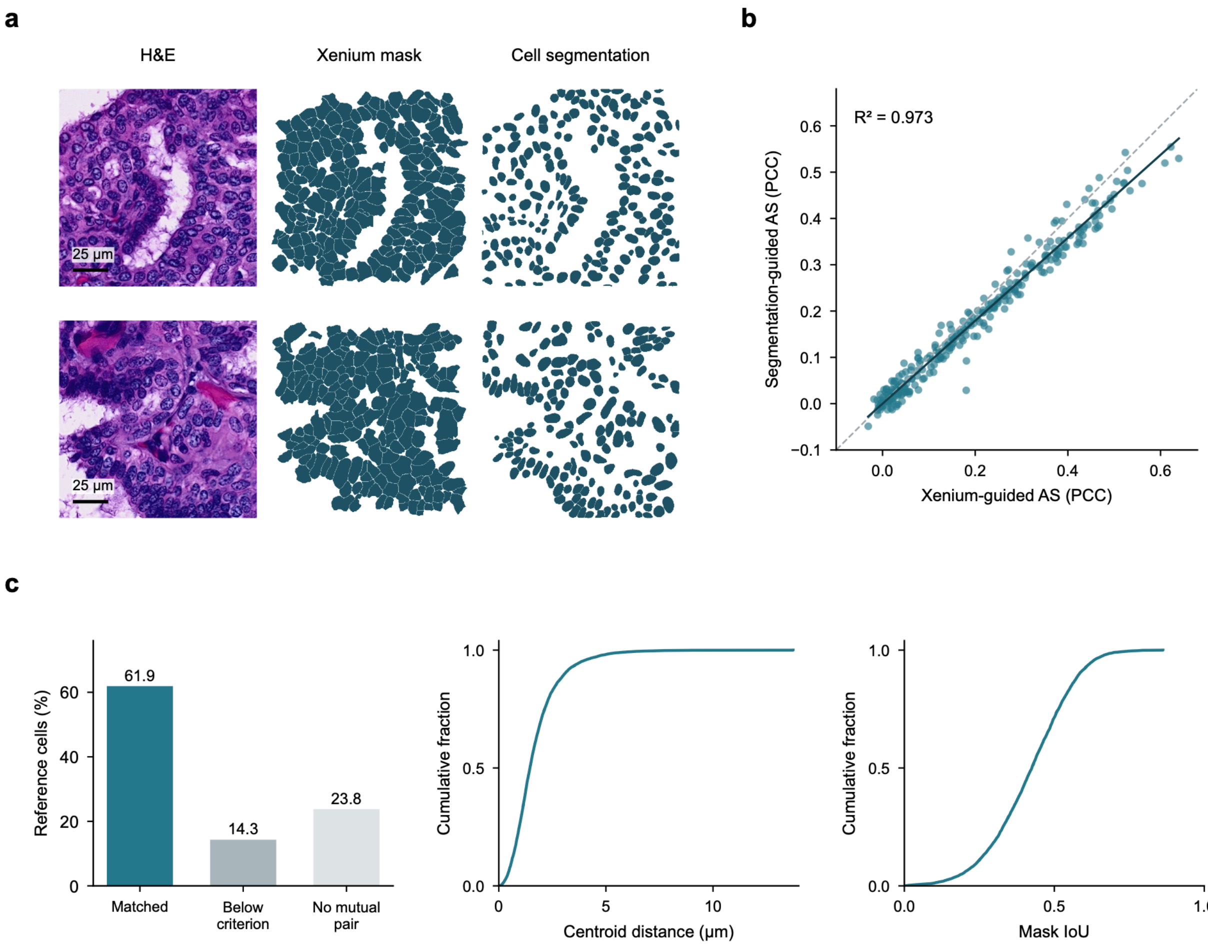


**Supplementary Fig. 4 | Reference-projected molecular representations across eight patients.**

UMAP reference projections for eight patients (rows), comparing Xenium, multi-tissue AS and nine comparator methods (columns). Analyses include 26,068 matched 55-µm positions and 280 genes. Within each patient, predictions undergo the same measured-expression-fitted standardization, principal component analysis and UMAP transformation. Colours retain the measured reference-state assignment of corresponding positions; plotting limits are shared across each row. All matched positions are shown without subsampling. C1–C8 are defined separately within each patient and do not imply equivalent identities between patients. The displayed n denotes spatial positions, not patients.

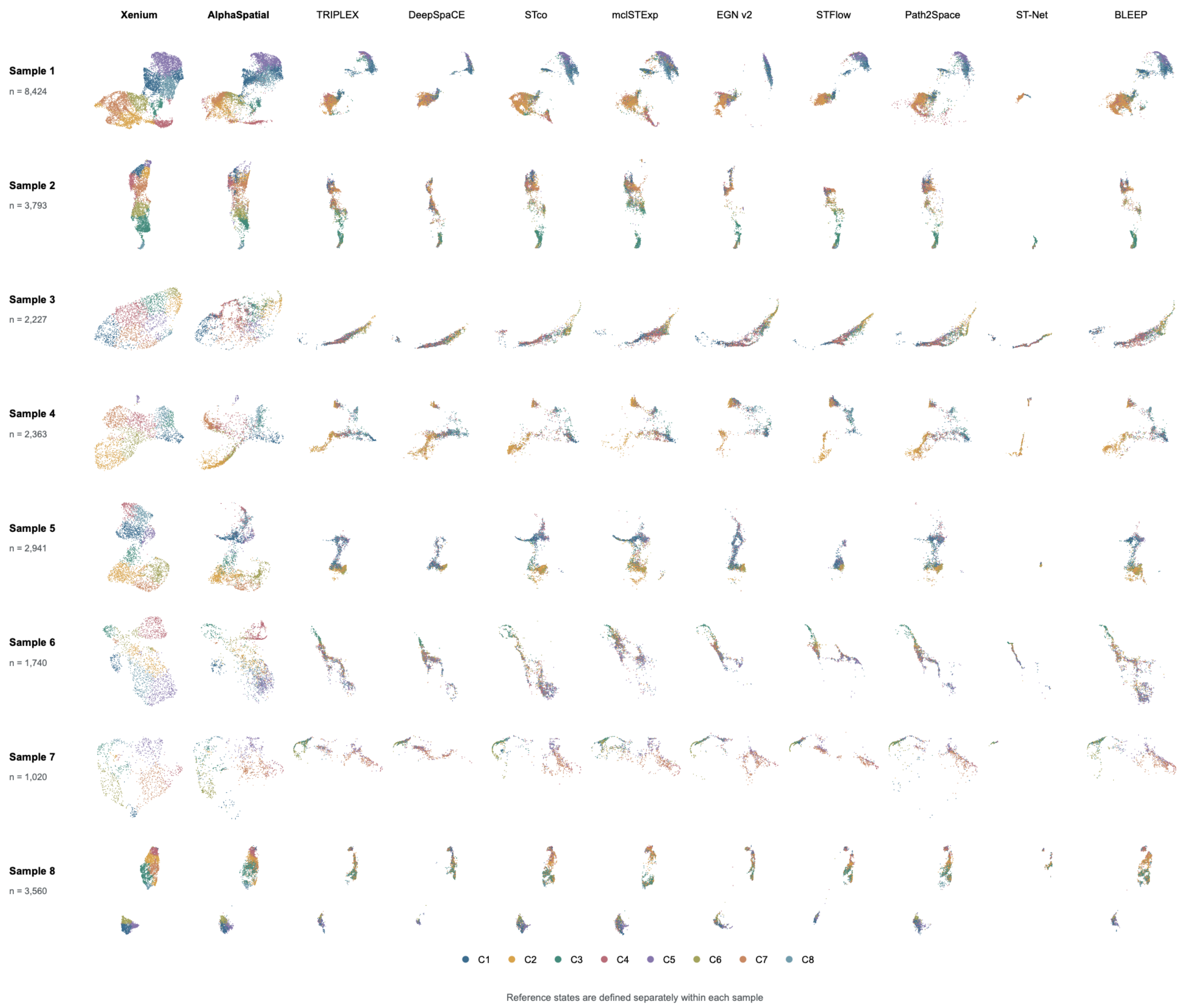


**Supplementary Fig. 5 | Spatial aggregation and multigene representation fidelity.**

a,b, Mean absolute error between measured and predicted gene–gene correlation matrices (a) and agreement between their 30-principal-component representations, assessed by linear centred kernel alignment (CKA; b), at single-cell and 55-µm resolution. c, Macro-F1 for assignment to eight reference-derived molecular states, defined separately at each scale. d, CKA for spatial bins versus occupancy-matched random groups. Random grouping preserved measured–predicted cell pairs and group sizes; results were averaged over ten randomizations per patient. e,f, CKA (e) and measured-expression variance retained after aggregation (f) across 27.5-, 55- and 110-µm bins. Variance retention denotes the median gene-wise ratio of bin-mean to cell-level variance. Analysis included 1,733,702 matched cells from eight independent patients. Bin expression was calculated by averaging absolute cell counts, without total-expression normalization. In a–d, connected points represent patients and horizontal bars indicate means; e,f show means ± s.d. Tests were two-sided exact paired Wilcoxon tests, with Holm correction across the two primary metrics separately for cell-versus-spatial and random-versus-spatial comparisons.

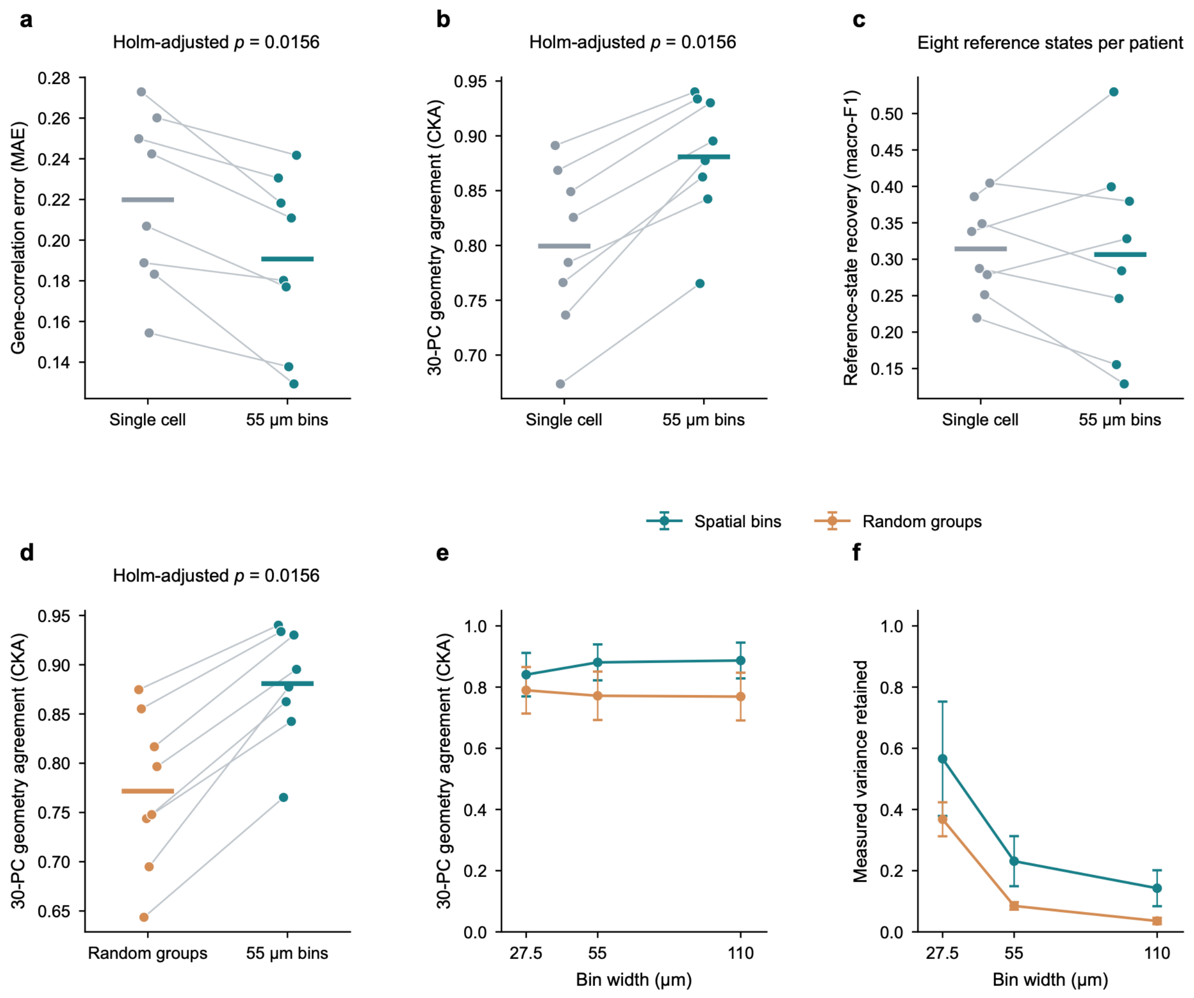

**Supplementary Fig. 6 | Spatial-state abundance, treatment response and regional composition.**

**a,** Abundance of four states as a percentage of epithelial-associated bins in 33 responders and 48 nonresponders. **b,** Spearman correlations between state abundance and residual invasive tumour size in 79 patients; error bars show 95% confidence intervals from 2,000 patient bootstraps. **c,** Within-patient differences in predicted cell-type proportions between S1 and other epithelial-associated regions in 80 patients. Differences are S1 minus other regions, in percentage points; unknown and tied assignments remain in the denominator. In a,c, points represent patients; boxes show medians and interquartile ranges, with whiskers extending to observations within 1.5 interquartile ranges. Tests used two-sided Welch tests (a) and 9,999 permutations for Spearman correlations (b), with Benjamini–Hochberg correction across four states separately for each endpoint. Panel c is descriptive; other regions are not normal tissue.

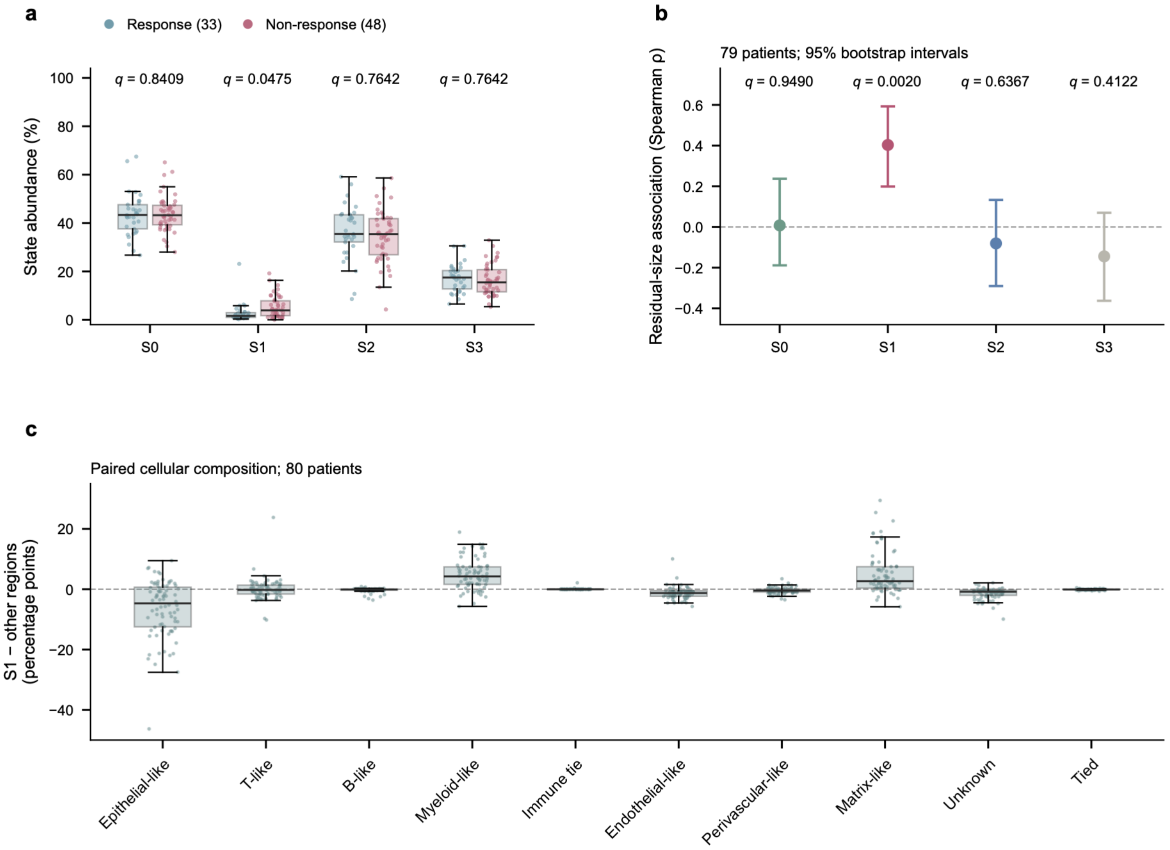

a
Response (33)
Non-response (48)
State abundance (%)
q = 0.8409
q = 0.0475
q = 0.7642
q = 0.7642
S0
S1
S2
S3
b
79 patients; 95% bootstrap intervals
Residual-size association (Spearman ρ)
q = 0.9490
q = 0.0020
q = 0.6367
q = 0.4122
c
Paired cellular composition; 80 patients
S1 − other regions (percentage points)
Epithelial-like
T-like
B-like
Myeloid-like
Immune tie
Endothelial-like
Perivascular-like
Matrix-like
Unknown
Tied
